# Immunocompetent murine models recapitulate the heterogeneous tumor immune microenvironment of human liposarcoma

**DOI:** 10.1101/2025.01.31.634916

**Authors:** Amanda M. Shafer, Emma Kenna, Lexi-Ann F. Golden, Ahmed M. Elhossiny, Kyle D. Perry, Jodi Wilkowski, Wei Yan, Brynn Kaczkofsky, J McGue, Scott C. Bresler, Adam H. Courtney, Jessie M. Dalman, Craig J. Galban, Wei Jiang, Carlos E. Espinoza, Rashmi Chugh, Matthew K. Iyer, Timothy L. Frankel, Marina Pasca di Magliano, Andrzej A. Dlugosz, Christina V. Angeles

**Author notes:** **Authorship note:** AMS, EK, LFG contributed equally to this work and are co-first authors. **Corresponding Author:** Christina V. Angeles, MD, FACS, FSSO, University of Michigan, Rogel Cancer Center, 1500 E. Medical Center Drive, 3304 Cancer Center, Ann Arbor, MI 48109.

## Abstract

Liposarcoma (LPS) is the most prevalent soft tissue sarcoma. The most common biological subtypes are well-differentiated (WDLPS), a low-grade disease that can evolve to high-grade dedifferentiated liposarcoma (DDLPS), with increased rates of recurrence and metastasis and low response rates to systemic therapies. Preclinical testing of immunotherapeutics for LPS has been held back by the lack of an immunocompetent mouse model. Here, we present an autochthonous immunocompetent LPS mouse model, ACPP, with targeted deletion of Trp53 and Pten in adipocytes to mimic signaling alterations observed in human LPS. Similar to humans, ACPP mice produce WDLPS, DDLPS, and tumors that exhibit both WD and DD components. Murine and human DDLPS tumors possess transcriptional similarities, including increased expression of oncogenes Cdk4 and Hmga2 and reduced expression of the tumor suppressor Cebpa; furthermore, both mouse and human DDLPS exhibit heterogenous T cell infiltration. Syngeneic cell lines derived from ACPP DDLPS reliably produce tumors following orthotopic implantation, each with distinct growth patterns, aggressiveness, and immune profiles. These unique models provide much needed tools to understand the complex immunobiology of LPS and greatly accelerate the pace of preclinical studies aimed at uncovering more effective new therapies for patients with this aggressive malignancy.

## INTRODUCTION

Liposarcoma (LPS) is the most common type of soft tissue sarcoma (STS) and arises in adipose tissues throughout the body. The most common biological types of liposarcoma are well-differentiated (WDLPS) and dedifferentiated liposarcoma (DDLPS). WDLPS, a low-grade disease, can evolve to DDLPS, a high-grade form that is associated with higher rates of local recurrence and distant metastasis (1, 2). WDLPS is treated with surgery alone and has a relatively good prognosis (5-year survival of 85%) (3); in contrast, patients with DDLPS have limited benefit from current systemic treatments. Surgical resection is the most effective treatment option for DDLPS, yet patients have a high risk of recurrence (4). Patients who recur with metastatic disease have poor overall survival of only 12-15 months due to modest response rates to cytotoxic chemotherapy (5, 6). The development of improved systemic therapies for LPS has been limited, leaving our first line therapy, doxorubicin-based chemotherapy, the same since the 1970s (7, 8). Reasons for the lack of progress in identifying more effective treatments include the inability to translate promising preclinical findings from *in vitro* and human xenograft models into effective patient treatments (9, 10).

While immune checkpoint inhibitor (ICI) immunotherapy has shown tremendous efficacy in many solid tumors, investigation of the immune biology of sarcoma has lagged behind research for other malignancies (11). Recently, however, sarcoma-focused immunology studies have demonstrated links between the tumor immune microenvironment (TIME) and patient outcomes (12-14). One of the first clinical trials, SARC028 (NCT02301039), a phase II, multicenter trial of the ICI pembrolizumab, showed promising activity in select histologic subtypes of advanced STS including DDLPS (15). However, there was only a 20% overall response rate. The causes underlying the limited efficacy of ICI in DDLPS are poorly understood. Limited data from human samples suggest that improved outcomes are associated with pre-existing immune responses (14, 15). The few DDLPS patients who responded to ICI in SARC028 had pre-treatment tumors with a higher density of effector memory cytotoxic CD8+ T cells (T_EM_) compared with non-responders (16). These tumors also contained tertiary lymphoid structures (TLS), an ectopic lymphoid structure with T-cell and B-cell rich regions (17). This suggests that productive immune responses occurred in at least a subset of patients. This is encouraging, but to understand what underlies this immune responsiveness, it is crucial to understand the LPS TIME.

Genetically engineered mouse models (GEMMs) are critical to preclinical immuno-oncology studies (18). Using GEMMs, we can study mechanisms of action in the context of relevant genetic drivers; furthermore, tumors in these models develop with a stroma and microenvironment that more dependably mimic human disease than cell-line derived models (19). Currently, immunotherapeutics for LPS remain untested *in vivo* due to the lack of a reliable immunocompetent LPS model. In this study we generated and validated a GEMM that carries key genetic aberrations to drive signaling alterations found in human WDLPS/DDLPS.

WDLPS and DDLPS are characterized by amplification of chromosome 12q13-15, which contains the major proto-oncogenes murine double minute 2 (*MDM2*) and cyclin-dependent kinase 4 (*CDK4*) (20). *MDM2* is a negative regulator of the tumor suppressor p53, and *CDK4* plays a crucial role in the control of cell cycle progression (9, 21, 22). While over 90% of WDLPS and DDLPS cases show copy number amplification of *MDM2* and ensuing increased proteins levels, *CDK4* expression is less consistent (23). Additionally, about 7% of human WDLPS/DDLPS have a *TP53* mutation instead of a *MDM2* amplification (20). While WDLPS and DDLPS are genomically similar, histopathological and clinical differences distinguish them. Human tumors frequently have both DDLPS and WDLPS components and are classified as DDLPS. Conditional inactivation of PTEN and p53 in mice with an adenovirus expressing Cre recombinase (Ad5CMV/Cre) leads to the development of multiple types of LPS (24). While PTEN mutations are not routinely found in human LPS (20), ∼80% of DDLPS have phosphorylated AKT (25). Knocking out *Pten* mimics the effect of constitutive phosphatidylinositol-3-kinase (PI3K)-AKT signaling. *MDM2* amplification is mimicked by knocking out its target operator p53.

To generate a reliable, spatially restricted LPS mouse model, we designed an adipocyte promoter-driven Cre fusion molecule and used an adenovirus-associated virus (AAV8) vector that elicits minimal immune response (26-28). Here, we characterize the autochthonous AAV8-**<u>A</u>**p2.2-eGFP/**<u>C</u>**re ***<u>P</u>****ten^f/f^;Tr**<u>p</u>**53^f/f^* C57BL/6 LPS model, hereafter called ACPP, including the histology, genomics, transcriptomics, and immune microenvironment of LPS tumors. We found both low grade WDLPS and high grade DDLPS that exhibited histologic features and transcriptomes similar to human LPS. Moreover, like human tumors, many DDLPS tumors arising in ACPP mice showed an associated WDLPS component. Strikingly, like a subset of human DDLPS tumors (16), 20% of ACPP DDLPS tumors had high tumor infiltrating lymphocytes (TILs). We created 3 syngeneic DDLPS cells lines of variable aggressiveness based on histological features and correlative growth rates, each with consistent orthotopic engraftment. These preclinical models provide an important tool for investigating the mechanisms of combination therapies and novel immuno-oncology treatment approaches for LPS.

## RESULTS

### Targeted deletion of Trp53 and Pten in adipocytes yields murine LPS histologically similar to human LPS

We designed a viral construct using an adipocyte-specific *adiponectin* promoter (mAP2.2) (26) to drive a GFP-Myc tag-Cre fusion molecule (29, 30) (Figure 1A). We packaged this construct into an AAV8 vector, which effectively transduces adipocytes and induces a negligible immune response (26-28). We confirmed transduction of adipocytes and Cre recombination after injecting AAV8-Ap2.2-eGFP/Cre (1×10^10^ vector genomes (vg) or 1×10^11^ vg) into the flank fat pad of *mT/mG* reporter mice, which express a constitutive membrane-targeted tdTomato transgene (mT) and STOP sequence that are excised by Cre recombinase, resulting in a switch to expression of membrane-targeted green fluorescent protein (mG) (Supplemental Figure 1). There was loss of mT and robust mG expression in adipocytes at the injection site. Although GFP fluorescence could be attributed to viral GFP/Cre expression, membrane localization coupled with loss of mT is consistent with successful Cre-mediated recombination to yield the expected reporter switch. We found sparse cells with mG expression in the adjacent interscapular fat pad and no expression in the lung, similar to control without injection (Supplemental Figure 1). These data are consistent with successful adipocyte-specific transduction, resulting in Cre-mediated recombination.

**Figure 1:**
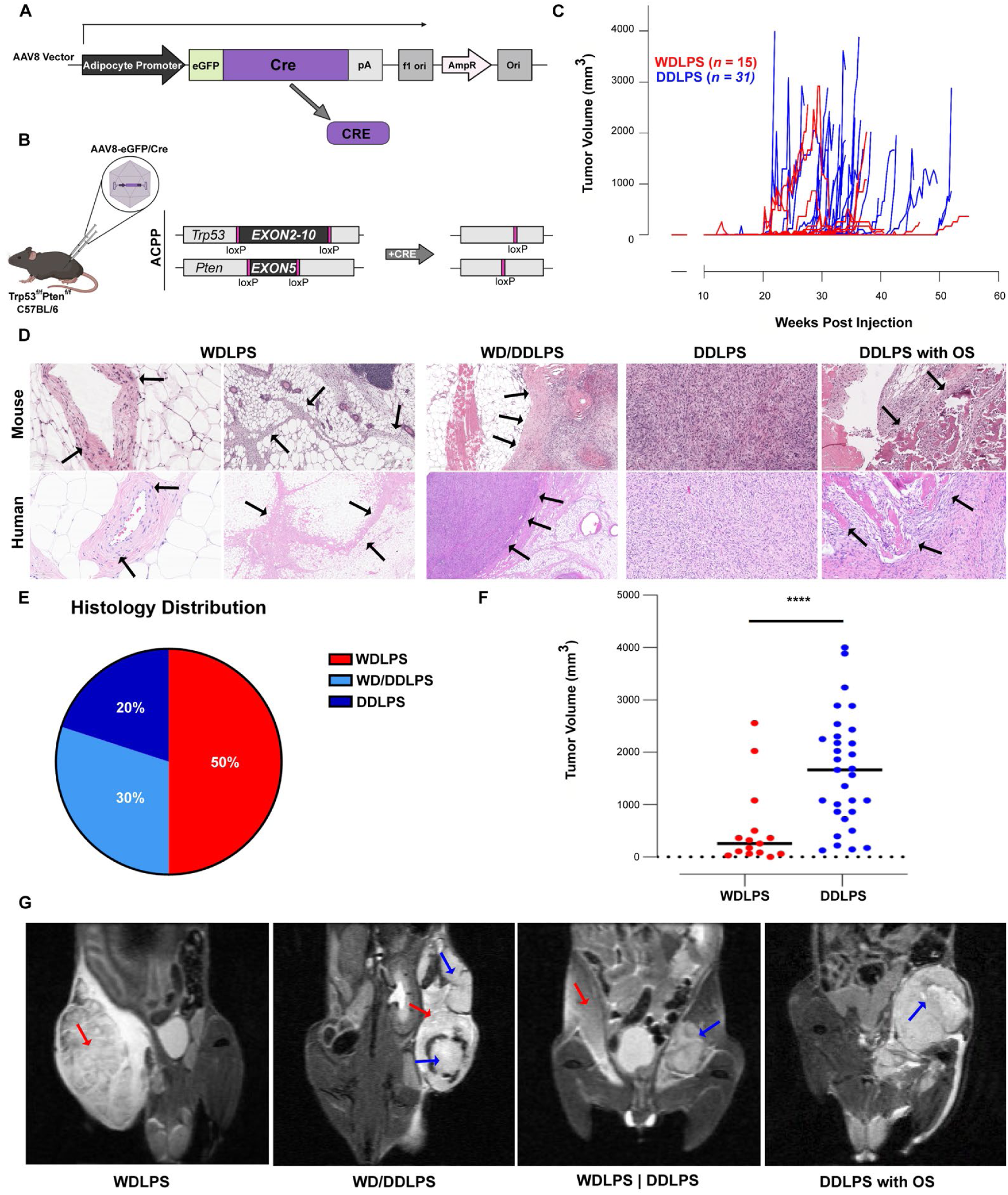
Targeted deletions of *Trp53* and *Pten* in ACPP mice generate LPS histologically similar to human LPS. (**A**) Creation of an adipocyte-specific promoter sequence to drive an eGFP-Myc tag-Cre fusion molecule. (**B**) Schematic of orthotopic injection of AAV8-Ap2.2-eGFP/Cre into *Trp53^f/f^Pten^f/f^* C57BL/6 GEMM, which generates autochthonous LPS tumors. (**C**) Tumor volume (mm^3^) growth curves in weeks post-injection of AAV8-Ap2.2-eGFP/Cre for WDLPS (red; *n* = 15) and DDLPS (blue; *n* = 31). (**D**) H&E of ACPP WDLPS, mixed DDLPS with WDLPS components, and DDLPS (top row) show similar histological findings as human subtype-specific LPS (bottom row). Features of murine and human tumors (left to right): WDLPS: atypical cells (arrows) entrapping vessels; WDLPS: fibrous septa (arrows) containing cells with atypical nuclei; WD/DDLPS: stark transition (arrows) between WDLPS and DDLPS components; DDLPS: consisting of atypical spindled and pleomorphic cells that exhibit a storiform to fascicular architecture; DDLPS: containing areas of bone formation (arrows), compatible with heterologous osteosarcomatous (OS) differentiation. (**E**) Histology distributions of ACPP LPS tumors. Red = WDLPS; Light blue = WDLPS/DDLPS; Dark blue = DDLPS. (**F**) Tumor volume (mm^3^) of WDLPS (red) and DDLPS (blue). Mean tumor volume is shown. Statistical differences determined by student’s t-test. Significance indicated by **** *P* < 0.0001. (**G**) MRI of APCC LPS tumors: WDLPS on right flank (far left), WDLPS transitioning into DDLPS on left flank (left), WDLPS on right flank and DDLPS on left flank (right), and DDLPS with OS differentiation on left flank (far right). Red arrows, WDLPS; blue arrows, DDLPS.

To replicate MDM2 amplification and p53 suppression observed in human LPS (20) and mimic the consistent activation of AKT in up to 80% of human LPS tumors (22, 25), we crossed C57BL/6J-congenic *Trp53^flox^*mice with C57BL/6J-congenic *Pten^flox^* mice to achieve *Trp53* and *Pten* knockout. Mice were crossed for multiple generations to establish homozygous *Trp53^f/f^Pten^f/f^* on a C57BL/6 background. We injected AAV8-Ap2.2-eGFP/Cre into bilateral flanks of *Trp53^f/f^Pten^f/f^*C57BL/6J mice, hereafter called ACPP (*n* = 53 total; *n* = 24 male and *n* = 29 female) (Figure 1B). LPS penetrance was 76% (80 LPS tumors/106 injections), without differences between sexes (77% in males and 74% in females; *P* = 0.7278). Some ACPP tumors developed around 3 months, and others during the subsequent 12 months post-injection (Figure 1C). We identified morphological features consistent with human WDLPS and DDLPS, including tumors with mixed WD and DD components seen in some human tumors (Figure 1D; see Supplemental Table 1 and Supplemental Figure 2, A and B). One DDLPS had a heterologous component of osteosarcomatous (OS) differentiation, analogous to a phenomenon sometimes seen in human DDLPS (Figure 1D, far right panel). Histological analysis of ACPP LPS tumors identified 50% WDLPS, 30% WDLPS/DDLPS, and 20% DDLPS (Figure 1E). DDLPS with WD components were grouped with DDLPS for further analysis following standard pathological classification of mixed tumors in patients (31). Since mice were injected in bilateral flank fat pads, some tumors were removed at a smaller size if they grew more slowly than the contralateral tumor, which reached the protocol’s maximum size endpoint. Therefore, there was a significant difference in the median size of histologically different tumors—288 mm^3^ for WDLPS and 1666 mm^3^ for DDLPS (Figure 1F; *P =* 0.0006)—suggesting that ACPP WDLPS could evolve to DDLPS over time, consistent with ongoing genomic instability seen in WDLPS/DDLPS human tumors (32). Immunohistochemical (IHC) staining of APCC DDLPS consistently demonstrated negligible levels of desmin expression, thereby excluding diagnosis of other high-grade myogenic tumors such as leiomyosarcoma (33) (Supplemental Figure 2, C and D). Tumors did not exhibit morphologic features of other liposarcoma variants, such as the myxoid stroma of myxoid liposarcoma or pleomorphic lipoblasts of pleomorphic liposarcoma (PLPS), which themselves do not exhibit an associated WDLPS component (34). Upon histological review, 26 ACPP fat pads could not definitively be diagnosed as LPS; hence, they were classified as ‘indeterminate’ lipomatous tumor and excluded from penetrance calculation and further analysis (Supplemental Figure 2D). All mice in this study reached humane endpoints prior to developing lung or liver metastases, examined both macro- and microscopically. Non-contrast magnetic resonance imaging (MRI) was used to identify autochthonous ACPP tumors before they were palpable, allowing us to track tumor growth (Figure 1G and Supplemental Figure 3). Therefore, MRI presents the opportunity to begin therapeutic intervention after confirming tumor initiation and growth, but prior to the tumor being palpable.

### Multi-omic characterization of ACPP tumors reveals similarities to human LPS

Human WDLPS and DDLPS are genomically characterized by an overall low tumor mutational burden. At the same time, they have many copy number alterations (CNAs) (32), the most prevalent being amplification of chromosome 12q13-15, which contains MDM2 and CDK4 (20, 35). To understand the biological processes that lead to tumor development in the ACPP model, we performed genomic and transcriptional profiling of murine tumors and compared them to human LPS. By whole exome sequencing (WES) of murine tumors (*n* =14), we detected a low tumor mutational burden and substantial CNAs, with higher frequency in DDLPS than WDLPS (Figure 2A), consistent with human LPS (31, 32). Chromosomes 3, 5, and 15 largely had copy number gains, whereas chromosomes 2, 7, and 14 had frequent copy number losses (Figure 2A). Notably, we observed significant copy number losses on murine chromosome 7, which contains CCAAT/enhancer binding protein alpha (*C/ebpa)*. This transcription factor is crucial for adipose tissue development; it also acts as a tumor suppressor and serves as a master regulator of adipogenic differentiation (36, 37). In human DDLPS, reduced expression of C/EBPA can result from hypermethylation or allelic loss and is associated with worse disease-specific survival (31). ACPP DDLPS also demonstrated copy number gains on murine chromosome 5, which contains the gene encoding cyclin-dependent kinase 6 (*Cdk6)*. CDK6 is key for cell cycle progression, and its overexpression in cancer is often linked to uncontrolled tumor growth (38). Interestingly, ACPP DDLPS tumors with the most copy number losses throughout chromosome 11 (N1019 and N1692), where the mouse *Trp53* gene is located, had the highest *Trp53* loss (Supplemental Figure 4A). Further analysis of CNAs across ACPP tumors revealed significant differences in fraction of genome altered (FGA) when samples were stratified by histological classification and amount of tumor infiltrating lymphocytes (TILs) in the TIME. WDLPS had a significantly lower FGA compared to DDLPS, with low TILs (WDLPS vs. DDLPS-low TILs; *P* = 0.017), in keeping with human LPS (31). DDLPS with low TILs had a significantly higher FGA compared to DDLPS with high TILs (*P =* 0.024) (Figure 2B; Supplemental Figure 4B); however, this is likely due to low tumor purity resulting from high immune infiltrate.

**Figure 2.**
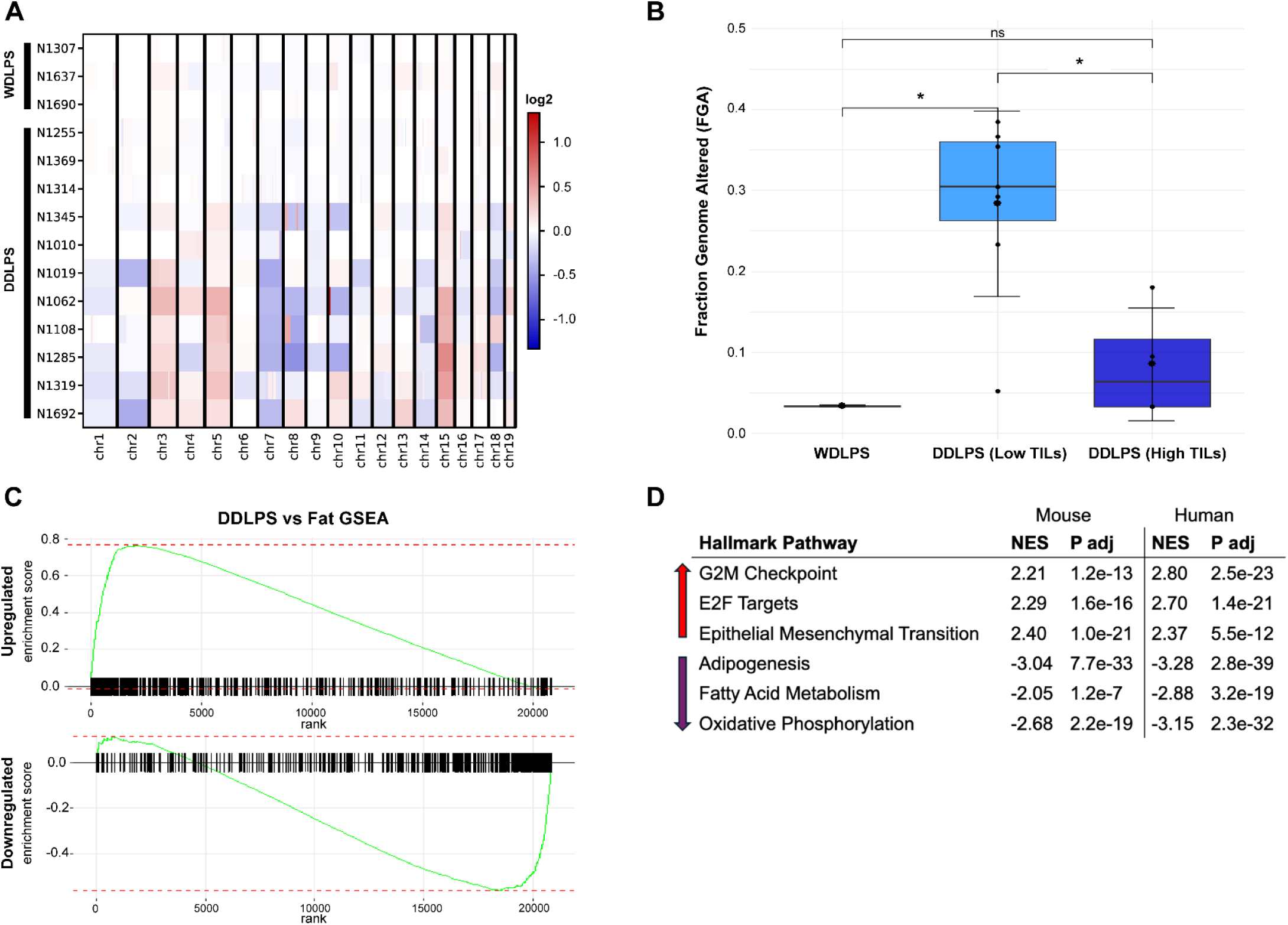
ACPP tumors transcriptomically resemble human LPS. (**A**) Heatmap of segmented log2-ratio read coverage comparing 14 murine ACPP tumors to normal fat using whole exome sequencing (WES) data, as analyzed by CNVkit. The heatmap represents genome-wide copy number alterations grouped by histological classification. Red blocks indicate chromosomal region gains; blue blocks indicate chromosomal region losses. (**B**) Boxplot depicting the Fraction of Genome Altered across distinct histologies (WDLPS *n* = 3, DDLPS (Low TILs) *n* = 7, DDLPS (High TILs) *n* = 4). Statistical differences determined by non-parametric Wilcoxon Rank Sum (Mann-Whitney) test. Significance indicated by * *P* < 0.05; ** *P* < 0.01. s (**C**) Gene set enrichment analysis (GSEA) of upregulated and downregulated gene sets in ACPP DDLPS (*n* = 10) compared to human DDLPS (*n* = 8) show concordance (NES 2.5, padj 2×10^-50^ and 2.5, 9.5×10^-35^, respectively) relative to species-specific normal fat, mouse *n* = 4; human *n* = 8. (**D**) Hallmark Pathway analysis of GSEA demonstrate similar up- and downregulated pathways in murine ACPP and human DDLPS.

Bulk RNA sequencing (RNA-seq) was conducted to determine gene expression profiles for murine LPS. Differential gene expression (DGE) analysis revealed that WDLPS (*n* = 3) and DDLPS (*n* = 10) had both differentially expressed up- and downregulated genes compared to normal fat controls (*n* = 4) (Supplemental Figure 4, C and D). We performed RNA-seq and DGE analysis on human tumors (WDLPS *n* = 8; DDLPS *n* = 8), relative to normal fat (*n* = 8). Gene set enrichment analysis (GSEA) showed that upregulated and downregulated gene sets in mouse ACPP DDLPS were similar to human DDLPS, with statistical significance (NES 2.5, padj 2×10^-50^ and 2.5, 9.5×10^-35^, respectively) (Figure 2C). We did not see the same concordance in WDLPS (Supplemental Figure 4E), likely due to it having more similarity to normal fat, with mostly upregulated and minimally downregulated gene differences (Supplemental Figure 4D). Additionally, MDM2, the most differentiated gene in human WDLPS compared to normal fat, was not amplified, likely because Trp53 is deleted in our model. Though the ACPP model is not designed to overexpress *Mdm2*, interestingly, both *Cdk4* and *Hmga2*—both found on chromosome 10 alongside *Mdm2*—were significantly overexpressed in DDLPS tumors (log2 FC 1.1, *P adj.* = 0.002 and 9.8, *P adj.* = 2.9×10^-9^, respectively) (Supplemental Figure 4C). In human DDLPS, *CDK4* and *HMGA2*, oncogenes on chromosome 12q with copy number amplification and overexpression, drive tumorigenesis (39). Under-expression of *C/EBPa* is important for DDLPS growth in xenograft models and is associated with worse patient survival (40-42). We also saw reduced expression of *C/epa* in murine ACPP DDLPS (Supplemental Figure 4C). Importantly, when comparing murine and human DDLPS against their respective species-specific normal fat, we identified concordance in significantly upregulated hallmark pathways involved with cellular proliferation (G2M checkpoint and E2F targets) and developmental programs (epithelial to mesenchymal transition) and in significantly downregulated pathways including the adipogenesis pathway and those involved with metabolic programs (fatty acid metabolism and oxidative phosphorylation) (Figure 2D). These findings highlight transcriptional similarities between human and murine DDLPS and support the ACPP model’s role as a valuable tool in preclinical applications.

*Autochthonous ACPP DDLPS tumors vary in density and composition of TILs.* All ACPP tumors (*n* = 80) were initially classified into low or high TIL status based on the abundance of lymphocytes and their proximity to tumor cells by H&E. High TILs were found in 18% of DDLPS tumors (Supplemental Table 1). The grade 2 DDLPS subset had 2.2-fold as many high TIL tumors as grade 3 (30% vs. 13%). DDLPS tumors without a WD component also had higher TIL than those with WD and DD components (25% vs. 13%). As expected, WDLPS lacked significant TILs, with none showing high TIL infiltrate (Supplemental Table 1). These findings recapitulate variability in immune infiltration seen in human LPS (16, 17). To characterize the TIME of ACPP tumors, we performed multiplex IHC (mIHC) analysis of immune cell populations, which revealed an overall low level of immune infiltrate in ACPP WDLPS tumors (Figure 3A) and immune heterogeneity in ACPP DDLPS tumors (Figure 3B-C). The mIHC panel probed for T cell markers CD3 and CD8, regulatory CD4 marker FOXP3, checkpoint marker PD-1, macrophage marker F4/80, and B cell marker CD19. Specified phenotype immune cells were quantified using InForm analysis (43-45). Murine DDLPS tumors were classified into low or high levels of TILs, hereafter called DDLPS^lo^ or DDLPS^hi^. DDLPS^hi^ had a percentage of CD3^+^ T cells out of total cells (determined by DAPI nuclear stain) twofold or greater than the median percentage of CD3^+^ for all samples; DDLPS^lo^ tumors had CD3^+^ percentages less than this threshold (*n* = 5 per classification). DDLPS^hi^ had an increase in percentage of CD3^+^ (Figure 3D), CD3^+^CD8^+^ (Figure 3E), and CD3^+^CD8^-^FOXP3^+^ Treg (Figure 3F) cells out of total cells in comparison to DDLPS^lo^ (*P* = 0.0079) and WDLPS (*P* = 0.0079). DDLPS^hi^ trended towards having more CD19^+^ B cells (Figure 3G) and fewer F4/80^+^ macrophages (Figure 3H) than DDLPS^lo^ (*P* = 0.55 and *P* = 0.55, respectively). Interestingly, while there was not a significant difference in the percent of macrophages between DDLPS^hi^ and DDLPS^lo^, each subtype produced distinct spatial patterns. Macrophages in DDLPS^lo^ were diffuse but confined to the periphery of DDLPS^hi^ (Supplemental Figure 5). Gene set variation analysis (GSVA) of bulk RNA-seq from ACPP LPS tumors (*n* = 13) and normal fat (*n* = 4) using defined signatures (46) to quantify immune and stromal cell populations showed that DDLPS tumors with high TILs (determined by H&E) had higher infiltration of T cells, memory B cells, and CD8 T cells compared to both DDLPS with low TILs and normal fat (Figure 3I). However, DDLPS with high TILs showed reduced macrophage infiltration (as determined by monocyte/macrophage enrichment) compared to DDLPS with low TILs but displayed higher macrophage infiltration relative to normal fat (Figure 3I). Altogether, these results emphasize the significant variability in immune cell infiltration within ACPP tumors, leading to distinctive TIMEs classified by immune cell population composition and unique spatial immune population patterns.

**Figure 3.**
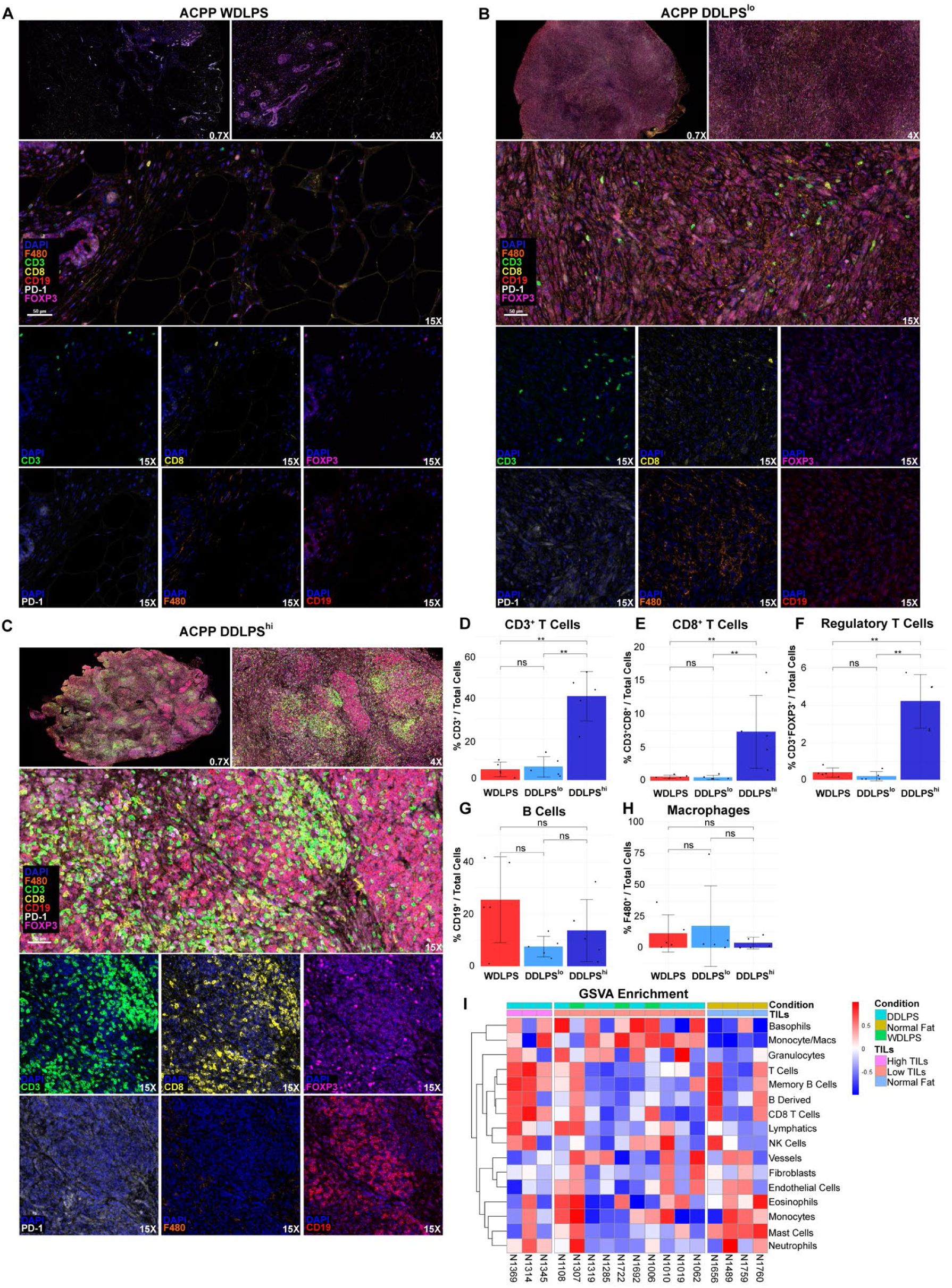
ACPP tumors have distinct TIMEs defined by variations in T cell populations. (**A-C**) Representative multiplex IHC (mIHC) of ACPP tumors representing WDLPS (**A**) and DDLPS with low (**B**) or high TILs (**C**). mIHC standard immune cell panel comprised of stains for CD3 (green), CD8 (yellow), FOXP3 (magenta), PD-1 (white), F4/80 (orange), and CD19 (red) with DAPI nuclear stain (Hoechst/blue). Shown at 0.7X, 4X, and 15X magnifications. Scale bar as shown. (**D-I**) Quantification of mIHC with inForm analysis for CD3^+^ (**D**), CD3^+^CD8^+^ (**E**), CD3+FOXP3^+^ (**F**), CD19^+^ (**G**), and F480^+^ (**H**) cells. DDLPS^hi^ were defined as those with CD3^+^ T cells ≥ 2x the cohort median; DDLPS^lo^ tumors had CD3⁺ levels below this threshold. Data, mean ± SEM; *n* = 5 samples/histological classification. Statistical differences determined by non-parametric Wilcoxon Rank Sum (Mann-Whitney) test. Significance indicated by * P < 0.05; ** P < 0.01. (**I**) Heatmap representing the value of the GSVA enrichment score, using mMCP gene signatures for immune cells, calculated for normal fat (*n* = 4) and ACPP tumor (*n* = 13) samples; grouped by H&E TILs classification.

### Intratumoral T cell phenotypes differ across histological subtypes of murine LPS

To further study T cell populations in APCC DDLPS tumors, we developed a targeted mIHC panel to investigate resident memory CD8+ T cell (T_RM_) populations. T_RM_ cells offer significantly stronger protective immunity in peripheral tissues compared to circulating central memory T cells (T_CM_) in both tumor and infectious disease models (47-49). Additionally, intratumoral T_RM_ cells are associated with better outcomes in patients with multiple solid malignancies (50-56). A core set of markers used to identify CD8^+^ T_RM_ include CD69, a marker of early T cell activation and tissue retention, CD103, an adhesion protein expressed by a subset of tissue memory CD8^+^ T cells, and PD-1, an inhibitory receptor expressed on CD8^+^ T_RM_ associated with prolonged antigen exposure in tumors and chronic infections (57-59). Using this panel, we investigated T cell heterogeneity in DDLPS (Figure 4, A and B). DDLPS^hi^ had an increase in CD3^+^CD8^-^ T cells classified as CD4^+^ T cells (Figure 4C) out of total cells in comparison to DDLPS^lo^ (*P* = 0.0095) and WDLPS (*P* = 0.057). DDLPS^hi^ also had an increase in CD8^+^ T cells compared to DDLPS^lo^ (*P* = 0.019) and WDLPS (0.057) (Figure 4D). CD8^+^PD1^+^ T cells were significantly reduced in DDLPS^hi^ versus DDLPS^lo^ (*P* = 0.025) (Figure 4F), while CD4^+^PD1^+^ T cells trended upward in DDLPS^lo^ versus DDLPS^hi^ (*P* = 0.07) (Figure 4E). CD8^+^ T cell subtype distribution by resident memory markers revealed distinct differences in the composition of CD8^+^ T cell phenotypes among LPS subtypes (Figure 4G; *P* < 2.2e^-16^). WDLPS was mostly comprised of CD69^+^ CD103^-^ PD1^-^ CD8^+^ T cells. CD8^+^ T cell phenotypes in DDLPS^lo^ were similar to those in WDLPS, with the largest proportion remaining CD69^+^ CD103^-^ PD1^-^, but with more CD69^+^ CD103^-^ PD1^+^ (T_RM_-like) and CD69^-^ CD103^-^ PD1^+^ CD8^+^ T cells (Figure 4G). CD8^+^ T cell phenotypes were considerably different in DDLPS^hi^ compared to DDLPS^lo^ and WDLPS; the largest T cell population in DDLPS^hi^, CD69^-^CD103^+^PD1^+^ CD8^+^, was not identified in other subtypes (Figure 4G). While CD69 is a marker of activation, it can be downregulated in chronic infection (60) and cancer (61). The proportion of CD69^-^CD103^+^PD1^-^ CD8^+^ T cells in DDLPS^hi^ was substantially increased, while proportions of CD69^+^CD103^-^ PD1^-^ CD8^+^, CD69^+^ CD103^-^ PD1^+^ (T_RM_-like) and CD69^-^ CD103^-^ PD1^+^ CD8^+^ T cells were decreased compared to WDLPS and DDLPS^lo^ (Figure 4G). While InForm analysis did not reveal any CD69^+^CD103^+^ CD8^+^ T_RM_ cells, they were identified by mIHC (Supplemental Figure 6). This was attributed to the low abundance of CD69^+^CD103^+^ CD8^+^ T_RM_ cells, which resulted in reduced likelihood of identification by InForm analysis. These data indicate that beyond the overall increased T cell infiltration in DDLPS^hi^, a notable variation in T cell phenotypes likely correlates with the transient, but not durable, anti-tumor response to immunotherapy observed in the few DDLPS patients from the SARC028 trial who had high intratumoral T cells (16). These findings support ACPP as a preclinical model for studying tumor-intrinsic and immune barriers to immunotherapy in LPS.

**Figure 4.**
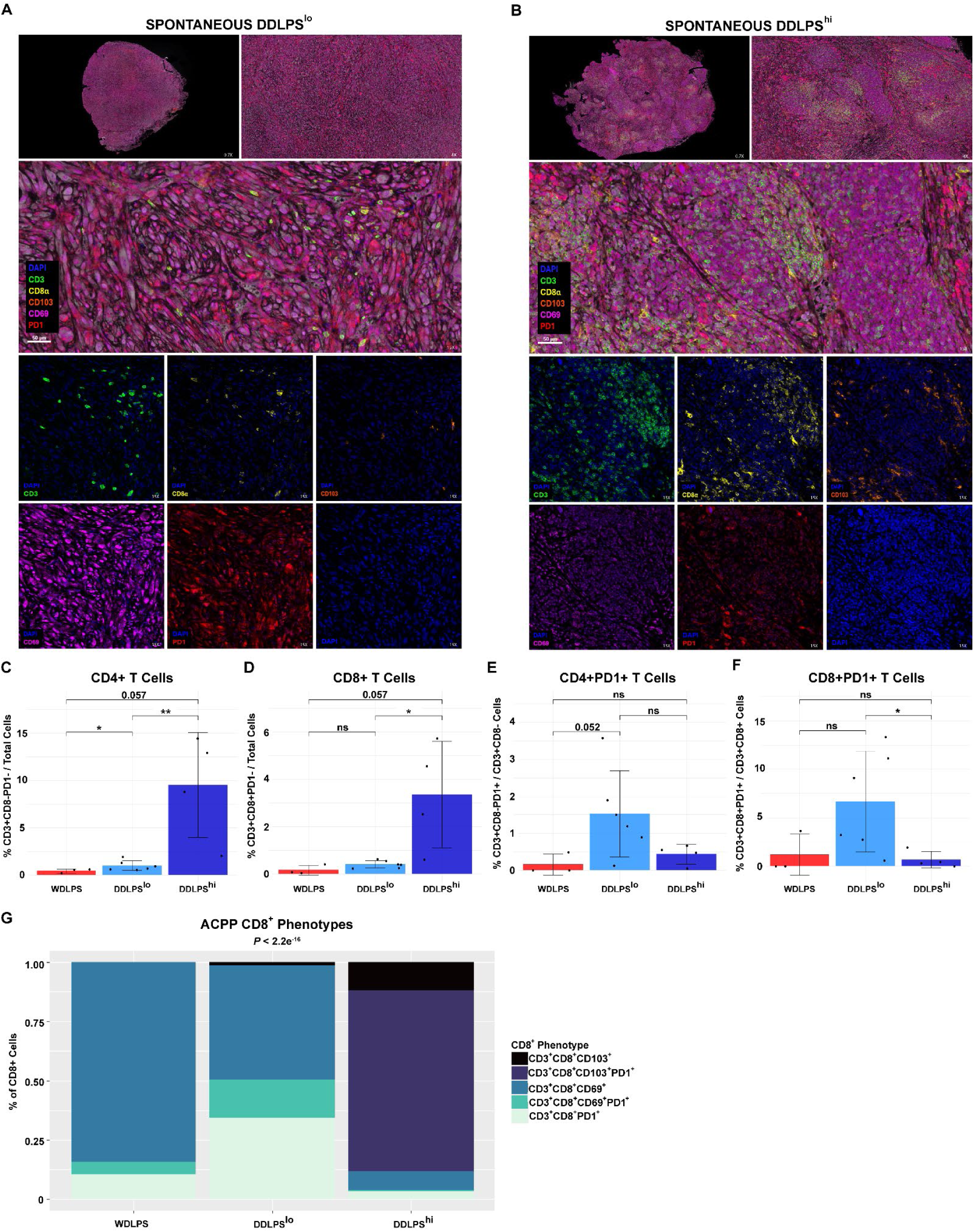
CD8+ T cell phenotypes differ between DDLPS^lo^ and DDLPS^hi^ tumors. (**A-B**) mIHC of ACPP tumors representing DDLPS with low (**A**) or high T cell density (**B**). mIHC resident memory T cell (T_RM_) panel comprised of stains for CD3 (green), CD8 (yellow), CD103 (orange), CD69 (magenta), and PD-1 (red) with DAPI nuclear stain (Hoechst/blue). Shown at 0.7X, 4X, and 15X magnifications. Scale bar as shown. (**C-F**) T cell quantification with inForm analysis for CD3^+^CD8^-^PD-1^-^ classified as CD4^+^ T cells (**C**), CD3^+^CD8^+^PD-1^-^ (**D**), CD3^+^CD8^-^PD-1^+^ (**E**), and CD3^+^CD8^+^PD-1^+^ (**F**) cells. DDLPS^hi^ were defined as those with CD3^+^ T cells ≥ 2x the median across all samples; DDLPS^lo^ tumors had CD3⁺ levels below this threshold. Data, mean ± SEM; *n* = 6 DDLPS^lo^ and *n* = 4 DDLPS^hi^ samples; *n* = 3 WDLPS samples. Statistical differences determined by non-parametric Wilcoxon Rank Sum (Mann-Whitney) test. Significance indicated by * *P* < 0.05; ** *P* < 0.01. (**G**) Stacked bar graph of the proportion of CD8^+^ phenotypes in ACPP tumors (*n* = 13) stratified by histological classifications; each phenotype is represented as a percentage of the whole; phenotypes whose percentage of the whole was 0 are not shown. Statistical difference in proportion composition determined by non-parametric Chi-square (Χ^2^) test.

### Syngeneic cell lines derived from autochthonous ACPP DDLPS form tumors following orthotopic injection

Syngeneic allografts are fundamental in translational research for their ability to rapidly and reliably produce tumors in immunocompetent mice. Hence, we generated and tested three syngeneic cell lines (N1011, N1018, N1343) originating from three distinct grade 3 DDLPS autochthonous tumors. Each cell line produced fully penetrant DDLPS allografts in syngeneic WT C57BL/6J mice (Supplemental Figure 7C). N1011 and N1018 were derived from ACPP DDLPS^lo^ tumors, whereas N1343 was from a DDLPS^hi^ tumor. The three cell lines showed distinct growth patterns *in vivo* when injected subcutaneously into the flanks of C57BL/6J mice (Figure 5A). N1018 allografts were the most aggressive and developed within 7 days of cell line injection. N1011 and N1343 were less aggressive and developed within 49-77 days post-injection. H&E analysis demonstrated morphological characteristics in cell line allografts similar to their respective autochthonous ACPP tumors (Figure 5B; Supplemental Figure 7A).

**Figure 5.**
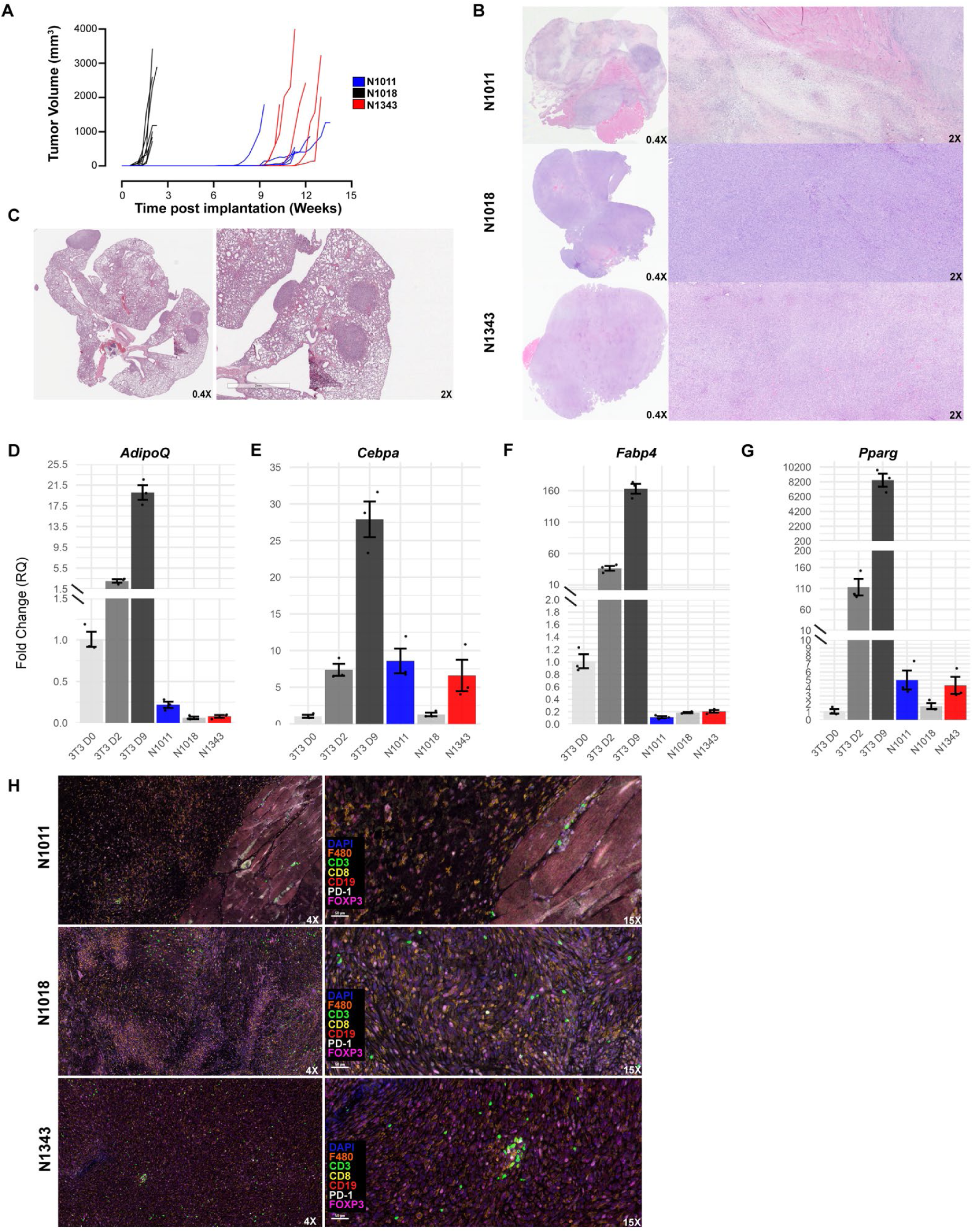
Syngeneic cell lines derived from autochthonous ACPP tumors develop orthotopic DDLPS tumors and lung metastases in WT C57BL/6 mice. (**A**) Tumor growth curves, represented by tumor volume (mm^3^), shown in weeks following cell line implantation for N1011 (blue; *n* = 5), N1018 (black; *n* = 7) and N1343 (red; *n* = 5). (**B**) Representative H&Es of orthotopic N1011, N1018, and N1343 tumors at 0.4X (left) and 2X (right) magnification. Scale bar as shown. (**C**) Representative H&E of lung metastasis developed after tail vein injected N1018. Scale bar as shown. (**D-G**) Analysis of gene expression levels in pre-adipocytes (3T3 day 0 = D0), adipocytes treated with differentiation factors for early differentiation (3T3 day 2 = D2) and late differentiation/mature adipocytes (3T3 day 9 = D9), and each cell line for adipocyte markers *AdipoQ* (**D**), *C/ebpa* (**E**), *Fabp4* (**F**), or *Pparg* (**G**). Gene expression was quantified using qRT-PCR and normalized to *Gapdh*. Samples were analyzed in triplicates. Data are presented as relative fold change (RQ) calculated by the 2A(-MCt) method. (**H**) Representative mlHC of orthotopic DDLPS (N1011 and N1018 *n* = 3, N1343 *n* = 2) stained with immune cell markers CD3 (green), CD8 (yellow), FOXP3 (magenta), PD-1 (white), F4/80 (orange), and CD19 (red) with DAPI nuclear counterstain (Hoechst/blue). Shown at 4X and 15X magnification.

### Tail vein injections of DDLPS cell lines results in lung metastasis with only the most aggressive N1018 cell line

Mice with orthotopic allografts did not develop visible or microscopic metastases prior to humane endpoint. To assess the metastatic potential of our DDLPS cell lines, WT C57BL/6J mice were injected intravenously through the tail vein for each line. Twenty-nine percent of mice injected with N1018 (*n* = 4 out of 14) developed experimental lung metastasis in an average of 36 days, while mice injected with N1011 or N1343 (*n* = 5 per group) had no evidence of lung metastasis over the course of 160 days. All lung tumors demonstrated the same histological characteristics of flank tumors, consistent with a diagnosis of grade 3 DDLPS lung metastasis (Figure 5C). This lung metastasis model reflects the predominant metastatic site in DDLPS (5, 6) patients, supporting its preclinical utility.

### Adipogenic differentiation gene expression varies among DDLPS cell lines

DDLPS cells express markers characteristic of an early stage of adipogenic differentiation (62). To determine the differentiation state of each syngeneic cell line, we compared the gene expression pattern of known adipocyte markers between cell lines and adipocytes in progressive stages of differentiation. Syngeneic cell lines showed very low expression of adipocytic markers *AdipoQ, Cebpa, Pparg,* and *Fabp4* compared to mature adipocytes (3T3 D9), as well as low levels of *AdipoQ, Pparg,* and *Fabp4* compared to early differentiated adipocytes (3T3 D2) (Figure 5, D-G). Notably, *Cebpa* expression varied across syngeneic cell lines. N1011 and N1343 showed *Cebpa* levels similar to those in early differentiated adipocytes (3T3 D2), while N1018 had significantly lower *Cebpa* expression, akin to pre-adipocytes (3T3 D0). Reduced expression in N1018 also coincided with the lowest levels of *AdipoQ*, suggesting less differentiation compared to N1011 and N1343 and correlating with the higher aggressiveness of N1018 (Figure 5, D and E).

### DDLPS allografts have variable TIL profiles

By mIHC immune cell analysis, we found distinct variations in the immune cell populations among the three DDLPS orthotopic allografts. N1011 had the highest proportion of macrophages and the lowest frequency of CD3^+^ T cells within the tumor, whereas N1018 had increased frequency of CD3^+^ T cells but retained the large, diffuse population of macrophages seen in N1011 (Figure 5H). N1343, which originated from an autochthonous DDLPS^hi^ tumor, had the largest proportion of CD3^+^ T cells within the tumor, with visual CD3^+^ T cell aggregates, and the lowest visual proportion of F4/80^+^ macrophages (Figure 5H). N1343 allografts did not share the same high TIL profile as the autochthonous DDLPS tumor it was derived from. This was not surprising, given that syngeneic cell line models typically do not form TLS, as cell lines may have evaded immunosurveillance in their original host and rapid growth typically does not allow time for TLS organization (19, 63-65). Nonetheless, the diverse TIL profiles among the three syngeneic cell line allografts provide a reliable model for generating DDLPS tumors in immunocompetent mice that reflect variable TIMEs seen in human tumors, thereby facilitating preclinical immunotherapy translational studies.

### Syngeneic cell line allografts have distinct differences in T cell phenotypes within the TIME

To characterize the TIME, immune populations from all three cell line allografts were analyzed by spectral flow cytometry (*n* = 5-10 per cell line). CD3^+^ T cell populations were clustered based on our T cell phenotype markers including CD4 and CD8, CD44 for antigen-experienced, CD69 for activation, CD103 and CXCR6 for residency, CD62L for circulation, and checkpoints LAG3 and PD-1 as markers of exhaustion (66). T-distributed stochastic neighbor embedding (t-SNE) analysis was used to compare CD3^+^ populations between cell line allografts (see Supplemental Figure 8A for pre-gating strategy). Using the FlowSOM clustering algorithm (67), we identified 15 CD3^+^ cell clusters, comprised of CD4^+^ T cells, CD8^+^ T cells, and CD4^-^/CD8^-^ double negative (DN) populations (Figure 6A). CD69, CD103, CXCR6, CD44, CD62L, PD-1, and LAG-3 markers were used to identify populations based on mean fluorescent intensities (MFI) that were mapped to the meta-clusters heat map generated by FlowSOM (Supplemental Figure 9A). We identified naïve T cells, characterized by elevated CD62L and lack of CD44 and CD69 (68); memory T cells, marked by shared expression of CD44 and CD69; resident-memory T cells (T_RM_) that expressed CD103 and/or CXCR6; exhausted T cells marked by dual expression of PD-1 and LAG-3 (66); and DN cells that were CD3^+^ but lacked CD8 and CD4 (69) (Supplemental Figure 9B). Most T cells in DDLPS allografts had a memory phenotype. t-SNE plots showed that N1343 allografts (derived from a DDLPS^hi^ ACPP tumor) harbored the most exhausted PD-1^+^/LAG-3^+^ CD4^+^ and CD8^+^ T cells and contained the least naive CD8^+^ T cells (Figure 6C). Specific T cell populations were also quantified by manual gating (see Supplemental Figure 8, B and C). Overall, N1343 allografts had the highest percentage of CD3^+^ T cells compared to N1018 (*P* = 0.023), with no significant differences between N1011 and N1018 (Figure 6D). N1018 had a higher proportion of CD4^+^ T cells compared to N1343 (*P* = 0.047) (Figure 6E) but a significantly lower proportion of CD8^+^ T cells (*P* = 0.00077) (Figure 6F). We did not find differences in the amount of PD-1^+^ or LAG-3^+^ CD8^+^ T cells between cell line allografts (Supplemental Figure 10, C and D). CD69^+^CD8^+^ T cells were similar in N1011 and N1018, but higher in N1343 compared to N1018 (*P* = 0.016) (Supplemental Figure 10E). CD103^+^ CD8^+^ T cells were more predominant in N1018 compared to N1343 and N1011 (*P* = 0.00028 and *P* = 0.069, respectively), while there were no differences between N1011 and N1343 (Supplemental Figure 9F). Dual expression of CD69 and CD103 is characteristic of tissue-resident memory (T_RM_) T cells, which, when present, are associated with improved response to immunotherapy (66, 70). The most aggressive cell line, N1018, had the highest percentage of CD69^+^CD103^+^ CD8^+^ T_RM_ compared to N1011 and N1343 (P = 0.028 and P = 6.8e-5, respectively) (Figure 6G), suggesting that even though N1018 allografts have overall lower CD8^+^ T cells than N1343, their tumor intrinsic properties may promote a more immunogenic TIME characterized by antitumoral T cell phenotypes, including T_RM_.

**Figure 6.**
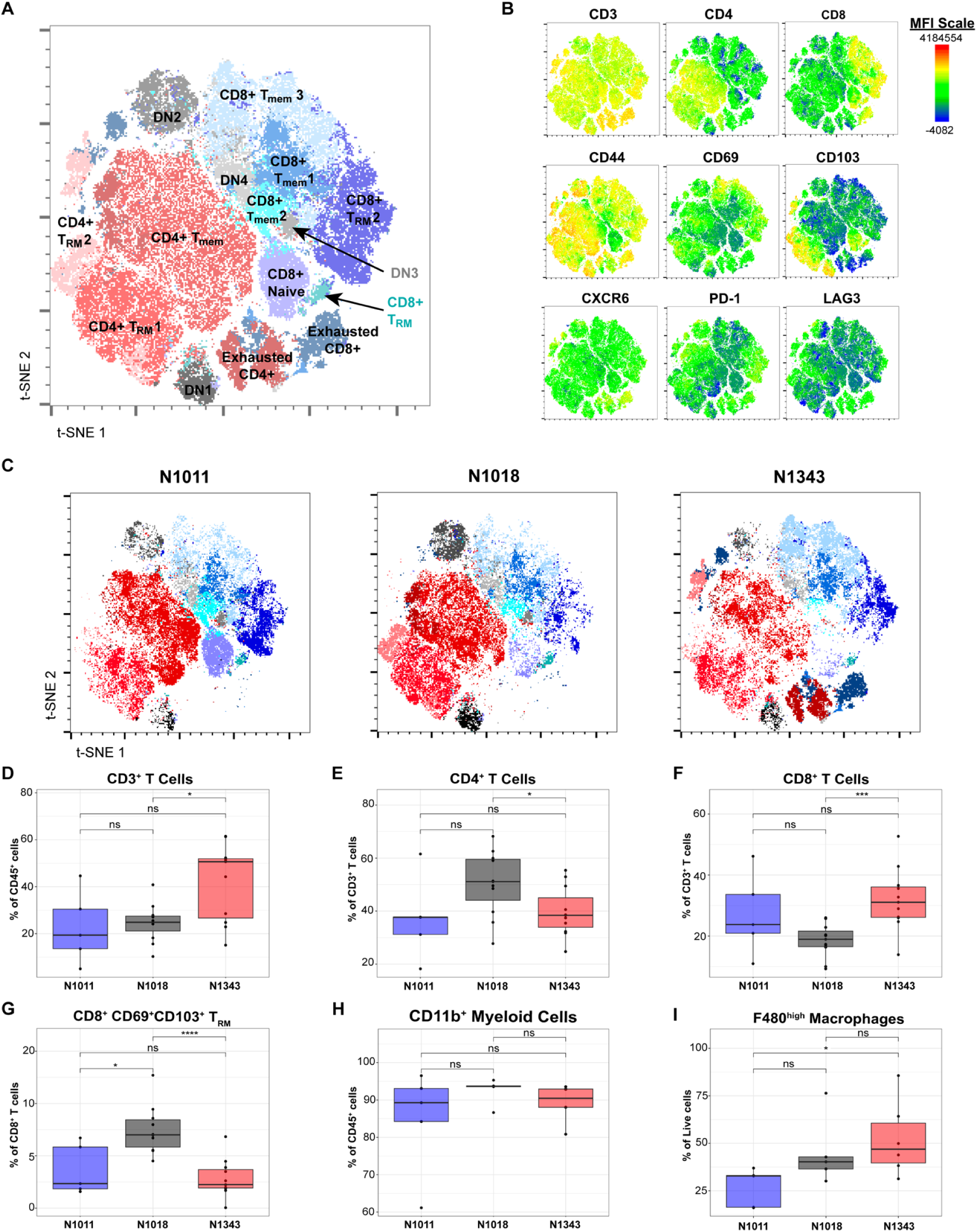
Orthotopic DDLPS have variable TIL profiles. (**A**) t-SNE map generated using FlowJo from spectral flow cytometry merged, pre-gated CD3+ cells from orthotopic N1011, N1018, and N1343 DDLPS in WT C57BL/6 mice (*n* = 3/cell line). (**B**) Heatmaps of markers (mean fluorescent intensity [MFI]) overlayed on t-SNE map of CD3^+^ T cells of all samples. MFI scale bar and fluorophores for each marker are shown. (**C**) t-SNE maps of independent orthotopic DDLPS tumors highlighting differences in clusters among cell lines. **(D-G)** Lymphocyte populations between orthotopic DDLPS tumors analyzed via spectral flow cytometry and manually pre-gated on lymphocytes/singlets/live/CD45^+^ (*n* = 5-10 per cell line). (**H-I**) Myeloid populations in tumors analyzed via spectral flow cytometry and manually pre-gated on all cells/singlets/live/CD45^+^ (*n* = 5-10 per cell line; see gating strategy in Supplemental Figure 8C). In (**D-I**), box plots represent mean (center line), length of the box represents 25th to 75th percentiles, and whiskers represent the 5th to 95th percentiles of data. Outliers are shown as dots outside of whiskers. *P* values were calculated by Kruskal-Wallis test. Significance indicated by * *P* < 0.05; *** *P* < 0.001; **** *P* < 0.0001. Flow cytometry gating strategy: see Supplemental Figure 8, A and B.

Myeloid populations, including CD11b^+^ myeloid cells and F4/80^+^ macrophages, play a critical role in modulating lymphocyte behavior in the TME (71). We found no differences in levels of CD11b^+^ myeloid cells (Figure 6H) and CD11b^+^F4/80^+^ macrophages (Supplemental Figure 10A) between cell line allografts. The percentage of F4/80^hi^-expressing macrophages, which is indicative of tumor-associated macrophages (TAMs), was highest in N1343 allografts compared to N1011 (*P* = 0.03) but showed similar levels to N1018 (Figure 6I). The percentage of CD19^+^ B cells was about two-fold higher in N1343 compared to N1018 and N1011 (*P* = 0.052 and *P* = 0.13, respectively) (Supplemental Figure 10B). Overall, these findings highlight the distinct immune landscapes present within each cell line allograft, with N1343 exhibiting a more exhausted T cell phenotype, and N1018 displaying a more immunogenic TIME characterized by a higher proportion of T_RM_ cells. Despite variations in T cell composition and phenotypes, myeloid and macrophage populations appeared relatively consistent across all three cell line allografts, suggesting that intrinsic DDLPS tumor properties may play a crucial role in shaping T cell immune responses. With their distinct characteristics, these cell lines used in concert provide a robust framework for investigating mechanisms of immune suppression, thereby laying the groundwork for developing future therapeutic strategies.

## DISCUSSION

To the best of our knowledge, this study describes the first autochthonous immunocompetent LPS mouse model that reliably mimics human LPS in several key aspects. First, histopathology shows that ACPP mice develop both WDLPS and DDLPS tumors, with some tumors exhibiting both WD and DD components, as is also seen in patients with LPS (Figure 1D; Supplemental Figure 2, A and B). DDLPS always arises in the context of WDLPS, thereby confirming that our model is WD/DDLPS, and not pleomorphic liposarcoma (31, 32). Second, ACPP DDLPS possesses transcriptional similarities with human DDLPS, including increased expression of oncogenes *Cdk4* and *Hmga2,* reduced expression of the tumor suppressor *Cebpa* (Supplemental Figure 4C), and concordance in the significantly upregulated and downregulated hallmark pathways (Figure 2, C and D). Third, and most compelling, is the heterogeneity in the TIME of the ACPP model, which develops tumors with variable T cell infiltration (Figure 3). In the correlative analyses of SARC028 clinical trial patients, the 20% of DDLPS patients who responded to anti-PD-1 had pre-treatment tumors with a higher density of cytotoxic tumor infiltrating T cells (16). The ACPP model generates about 18% of DDLPS^hi^ tumors, strikingly similar to the 20% of patients with high T cell tumor infiltrates (16). The autochthonous ACPP model offers a powerful research tool to investigate mechanisms underlying immune evasion in DDLPS, with the long-term goal to improve immunotherapy approaches for this disease.

Our group and others have shown that the presence of tissue-resident memory T cells (T_RM_) in the tumor microenvironment plays a significant role in the anti-tumor immune response and predicts better outcomes in patients with multiple solid malignancies (50-56). We found a striking difference in T cell phenotypes in DDLPS^hi^ compared to DDLPS^lo^ murine tumors. DDLPS^hi^ had a higher percentage of CD69^-^CD103^+^PD-1^+^ CD8^+^ T cells, indicating a T_RM_-like population that may have initially expressed CD69 during early activation but subsequently downregulated it, similar to patterns observed in chronic infection and selected solid tumors (60, 61). We identified only a few CD103^+^CD69^+^ CD8^+^ T_RM_ cells in DDLPS^hi^ tumors and none in DDLPS^lo^, suggesting that this is not a predominant T cell phenotype in liposarcoma (Figure 4G, Supplemental Figure 6). Overall, our data emphasize differences not only in the presence and density of T cells in LPS, but also in the specific phenotypic profiles of CD8^+^ T cells necessary for antitumor immunity. Our model can serve as a platform to evaluate new immunotherapy strategies and determine how LPS tumor immune cell populations affect treatment outcomes.

Analysis of pre-treatment biopsies from patients with DDLPS who had improved progression-free survival in phase II ICI trials revealed the presence of TLS (14, 15), an ectopic lymphoid structure with T-cell and B-cell rich regions and other immune populations (17). Our data suggest that DDLPS^hi^ tumors likely contain immature TLS, based on observed disorganized aggregates of T and B cells, as well as an increased abundance of exhausted T cells (Figure 3C) (72). This lack of functional immune activity can have different causes, including intrinsic T cell dysfunction or accumulation of immune suppressive myeloid cells (73). Pathological data from over 1,200 sarcoma patients with over 27 histological subtypes, including 62 DDLPS patients, demonstrated infiltration of predominantly pro-tumorigenic tumor-associated macrophages with density that exceeded that of TILs (74, 75). In pancreatic cancer, another “cold,” poorly immunogenic tumor, immunotherapy has largely been ineffective; however, studies of exceptional survivors showed the presence of memory T cells expressing T-cell receptors (TCRs) that recognize tumor antigens, even decades after the primary tumor was resected (76). Thus, even in poorly immunogenic tumors, if the immune system is engaged, long-term immune control of tumors is possible. Developing immunotherapeutic approaches for LPS patients will depend on further investigation of DDLPS tumors using both spatial and single-cell RNA and TCR sequencing, to deepen our understanding of immune interactions occurring in high TIL tumors.

We developed three syngeneic cell lines from three independent ACPP-derived DDLPS tumors that demonstrate 100% penetrance as orthotopic allografts in C57BL/6 mice. Syngeneic models are important research tools that develop tumors faster and are less expensive than spontaneous GEMM models; moreover, each cell line enhances reproducibility and exhibits specific traits that allow us to study tumor heterogeneity. In contrast to xenograft models used in LPS preclinical studies (42), syngeneic cell line *in vivo* models have an intact immune system, enabling study of the TIME and immunotherapies. Each ACPP-derived cell line described herein has distinct growth patterns, aggressiveness and TIL profiles. The most aggressive DDLPS line (N1018) is characterized by fast growth, metastatic potential, and the most dedifferentiated phenotype (lowest expression of *Cebpa*, *Pparg*, and *AdipoQ*) (Figure 5), but interestingly, has the highest population of CD8^+^ T_RM_ cells (Figure 6). This suggests a TIME that is potentially responsive to immunotherapy; however, the increased abundance of macrophages and other barriers may need to be overcome to prompt a durable response. This highlights the utility of our models for investigating tumor cell-immune interactions in response to chemotherapy, immunotherapy, and combination therapies in DDLPS cell line allografts in preclinical translational studies.

While ACPP mice and ACPP-derived DDLPS cell line allografts carry key genetic aberrations found in human LPS, a potential limitation is that these models do not have overexpression of MDM2 found in the majority of human WDLPS and DDLPS tumors, but do have p53 loss seen in a small fraction of patients (20). Despite promising preclinical results with MDM2 inhibitors *in vitro* and in patient tumor-derived DDLPS xenografts, including complete tumor regression in some models (10, 77-79), multiple clinical trials have not demonstrated a clear benefit in DDLPS patients (9). This outcome emphasizes the need for improved preclinical models, such as an adipocyte-specific MDM2-overexpressing GEMM that we are currently evaluating. Although preclinical animal models have limitations, testing potential new therapies requires leveraging a variety of models. Our ACPP mice and cell lines will allow exploration of the underlying mechanisms affecting how LPS tumor cells and the immune microenvironment interact in response to therapy. This approach is especially critical to advancing our understanding of and achieving breakthroughs in rare cancers like liposarcoma.

In conclusion, our new, highly penetrant immunocompetent GEMM for autochthonous LPS, along with cell lines that grow as allografts in syngeneic mice, provide much-needed immunocompetent animal models of this aggressive malignancy. These models closely mimic human disease and hold the potential to greatly accelerate the pace of uncovering new therapies to improve outcomes for patients with LPS.

## METHODS

Additional details can be found in Supplementary Materials.

### Sex as a biological variable

Male and female mice were used for all experiments. Data show similar findings for both sexes.

### Generation of AAV8-Ap2.2-eGFP/Cre Virus

The AAV8-Ap2.2-eGFP/Cre virus was made by the University of Michigan (U-M) Vector Core (Ann Arbor, MI). In brief, the AP2.2 gene block was synthesized by IDT. The eGFP-Myc-Cre gene block was PCR amplified from addgene plasmid #49056. The two fragments were assembled in a 3-part Gibson assembly into BstAPI/Xhol digested pCW3SL adeno-associated transfer plasmid (80) using NEB Gibson Assembly Master Mix following manufacturer’s instructions and transformed into NEB stable competent cells. Individual clones were picked and grown overnight before DNA miniprep. Assembly of the Ap2.2-eGFP/Cre plasmid was verified by Sanger sequencing. The plasmid was used to generate recombinant AAV8 virus by transfection of 293T cells and purified by iodixanol gradients. Titers of the viral preparations were determined by qPCR using a published protocol (https://www.addgene.org/protocols/aav-titration-qpcr-using-sybr-green-technology/).

### Mice

Mice were housed at and cared for by the U-M Unit for Laboratory Animal Medicine (ULAM) and the Angeles Lab. Animals were housed in standardized conditions to avoid intergroup variation.

### Trp53^f/f^Pten^f/f^ GEMM

Trp53^f/f^ mice (B6.129P2-*Trp53^tm1Brn^*/J, Jackson Laboratory, stock #008462) were crossed with Pten^f/f^ mice (B6.129S4-*Pten^tm1Hwu^*/J, Jackson Laboratory, stock #006440) to produce homozygous *Trp53^f/f^ Pten^f/f^* C57BL/6J mice. Genotyping was performed by TransnetYX (Cordova, TN).

### Viral Injections

AAV8-Ap2.2-eGFP/Cre virus was diluted with PBS to a low (2.3 × 10^10^ vector genomes [vg]) and high dose (2.3 × 10^11^ vg) and injected subcutaneously into the left and right flank fat pads of mT/mG indicator mice to assess Cre-mediated recombination in tissue. AAV8-Ap2.2-eGFP/Cre virus was diluted with PBS to concentrations ranging from 1.15 × 10^10^ vg to 1.62 × 10^12^ vg and injected subcutaneously into the left and right flank fat pads of *Trp53^f/f^Pten^f/f^* mice or wild type C57BL/6J mice as control. Mice were anesthetized with 2.5% isoflurane for all virus injections.

### Syngeneic cell line generation and culturing

Three cell lines (N1011, N1018, N1343) were generated from three independent autochthonous ACPP DDLPS tumors (see Supplemental Figure 6C). In brief, 500 mg of tumor were rinsed with Hanks’ Balanced Salt Solution and minced prior to digestion in collagenase IV solution [0.5 mg/mL DNAse (Sigma, 10104159001), 1 tablet Complete Mini EDTA-free Protease Inhibitor cocktail (Sigma, 11836170001), 5 mg/mL Collagenase Type IV (Sigma C5138-500mg)] for 30 minutes in a 37°C shaking incubator. Fetal bovine serum (FBS) was added to the suspension to stop the reaction. Samples were filtered through a 100 μM nylon mesh filter and pelleted via centrifugation at 450g and supernatant was discarded. Samples were resuspended in 1% FBS/PBS and filtered through a 40 μM nylon mesh filter. Samples were pelleted again as described above prior to being resuspended in growth media (GM) [DMEM HG F12 (Gibco 11320), 10% FBS, 100 U/mL Penicillin/Streptomycin, 2 mM L-glutamine, 2.5 µg/mL plasmoscin] and plated in a 6-well plate. Cells were cultured at 37°C in a humidified incubator with 5% CO_2_ and passaged when they approached confluency. Aliquots of cell lines were frozen down for storage at various passages (P3-P7) in 10% DMSO/FBS.

### Syngeneic cell line injections

For orthotopic allograft formation, 250,000-1,000,000 tumor cells were suspended in GM and mixed 1:1 in Matrigel (Fishers, 356231) to in 200 μL for subcutaneous injections. To assess metastatic potential, 100,000 tumor cells were resuspended in GM in 100 μL for tail vein injections. Cells were subcutaneously (flank) or intravenously (tail vein) injected into 8 – 12-week-old WT C57BL/6J mice (Jackson Laboratory, stock #000664). Mice were anesthetized with 2.5% isoflurane for all cell injections.

### Immunohistochemistry (IHC)

Tissues were fixed for 24 hours in 10% neutral-buffered formalin and transferred to 70% ethanol before being embedded in paraffin and sectioned. Hematoxylin (Leica, Surgipath, Cat #3801562) and eosin (Leica, Surgipath, Cat #3801601) (H&E) stain and desmin were performed on FFPE sectioned mouse tissues following standard protocol by the U-M Rogel Cancer Center Histology Core on the Ventana Discovery Ultra (Roche, Indianapolis, IN). See Supplemental Table 3 for antibody details. DAPI (Abcam, AB228549) was used to stain nuclei. mT/mG indicator mice samples were stained for GFP and tdTomato using the ABC method (81) to assess Cre-mediated recombination in adipocytes/fat pads and distant organs.

### Histopathological analysis

Histopathologic categorization was performed and independently categorized by two soft tissue pathologists using commonly accepted morphologic criteria for human WD/DDLPS. Diagnostic features of WDLPS included fibrous septa containing atypical mesenchymal cells with enlarged, irregular or hyperchromatic nuclei in the background of adipocytes. Typical identifying features of DDLPS included abrupt loss of adipocytic morphology, in which the tumor was composed of sheets of pleomorphic or highly atypical, spindled cells. During microscopic examination, the amount of tumor infiltrating lymphocytes (TIL) was approximated. Tumors with occasional lymphocytes in contact with tumor cells over ten fields of high magnification (400×) were designated low TIL; those with lymphocytes consistently in contact with tumor cells at medium magnification (200×) were designated low-moderate TIL. In high TIL tumors, lymphocytes were easily apparent at low magnification (100×) and at least focally obscured by clusters of sarcoma cells.

### Spectral Flow Cytometric Analysis

Unmixed FCS files were analyzed using traditional gating methods for lymphocytes (Supplemental Figure 7B) and myeloid populations (Supplemental Figure 7C). Three cell line allografts containing median levels of CD3+ T cells as determined via manual gating for each line were selected for dimensionality reduction analysis via t-distributed Stochastic Neighbor Embedding (t-SNE) using FlowJo^TM^ v10 software. Samples were pre-gated using the gating strategy in Supplemental Figure 7A and down sampled to 15K events per sample using DownSample v3.3.1. FCS files for each sample were concatenated prior to running t-SNE using the following settings: opt-SNE learning configurations at 1000 iterations, 30 perplexity, the random projection forest KNN algorithm, and the Barnes-Hut gradient algorithm (82, 83). Clustering was performed via FlowSOM v4.1.0. Multiple iterations were performed to determine the optimal number of clusters (16 populations were selected). Only clusters with > 500 cells were considered (*n* = 1 cluster excluded from analysis). The generated meta-cluster heatmap was used to name clusters (Supplemental Figure 9A).

### Multiplex fluorescent immunohistochemistry (mIHC) staining and imaging

5 µm paraffin-embedded tissue slices were placed onto charged slides, baked for 1 hour at 60°C, rehydrated as prior, and fixed with formalin. Multiplex staining was conducted (43) utilizing the OPAL platform and 6-plex manual detection kit (Akoya, #NEL861001KT). Briefly, slides were prepared for each round of staining using an antigen retrieval buffer with pH 9 or pH 6 (AR9 and AR6, Akoya Biosciences), followed by antibody application (see Supplemental Table 2 for antibody information) secondary antibody application, and fluorescent tyramide signal amplification. Slides were counterstained with 4’6-diamidino-2-phenylindole (DAPI) and a composite image obtained on the Vectra Polaris microscope. Ten regions of interest were identified for each tumor or slide; using InForm software (Akoya Biosciences), each cell was assigned a cellular phenotype and unique spatial location, allowing for quantification of proportion of tumor-associated immune cell subtypes, inter-cellular distances, and nearest neighbor calculations (43).

### Whole Exome Sequencing Analysis

Samples were aligned to mm39 using Burrows-Wheeler aligner. We used “CNVKit” for copy number variation analysis. We used “KaryoploteR” R package for visualizing coverage and log ratio. The log ratio for *Trp53* and *Pten* exon 5 was calculated using a weighted average of CNVKit bin-level calls of bins overlapping gene regions. The fraction of genome altered (FGA) was determined by dividing the total length of altered segments by the mouse genome size (∼2.7 Gb).

### Bioinformatics Analysis

Mean tumor volume (mm^3^) analysis and visualization was performed using GraphPad Prism 10 (GraphPad Software, Inc). WES analysis was performed using the Python library CNVKit to detect copy number variants and alterations genome-wide from high-throughput sequencing data. Spectral flow cytometric and mIHC quantification data and FGA data were visualized using “ggplot2” to create boxplots and barplots and “pheatmap” to make heatmaps; “ggpubr” and “BayesFactor” were utilized to determine statistical significance between groups and proportion composition among groups, respectively. RNA sequencing count data were analyzed using DESeq2. Variance stabilizing transformation (vst()) function was used to generate normalized count matrix for visualization. PCAtools R packages were used to generate PCA plots. Differentially expressed genes between WDLPS vs. normal fat and DDLPS vs. normal fat were defined using DESeq() function with ‘condition’ and ‘batch’ as covariates. Gene Set Enrichment Analysis (GSEA) was done using the fgsea R package using the MSigDB Hallmarks (MH) gene set, which was downloaded using msigdb R package. GSEA was used to score enrichment of gene sets differentially expressed human in WDLPS vs. normal fat and DDLPS vs. normal fat in mouse RNAseq comparisons. The convert_human_to_mouse_symbols() function from nichenetR R package was used to convert human genes to mouse homologs. Gene Set Variation Analysis (GSVA) (84) was utilized to score cell-type signatures defined by murine Microenvironment Cell Population counter (mMCP) (46). Heatmaps visualizing expression were generated using “ggplot2” and “pheatmap” R packages. Volcano plots visualizing differential gene expression, calculated using DESeq() and lfcShrink(), were created using the “EnhancedVolcano” R package.

### Human sample collection

Human studies were performed in accordance with ethical regulations and pre-approved by the U-M Medical School Institutional Review Board (IRBMED) (#HUM00196054). Patients with soft tissue tumors and a pathological diagnosis of liposarcoma provided written informed consent to obtain fresh tissue from surgical specimens as part of the Sarcoma Research Working Group Biobank at the U-M Rogel Cancer Center.

### Study Approval

Animal studies were reviewed and approved by U-M IACUC under animal protocol #PRO00011576. Animals were housed, maintained, and handled according to guidelines in place at U-M, which is accredited by the American Association for the Assessment and Accreditation of Laboratory Animal Care (AALAC). All animal experiments complied with AALAC guidelines, US Department of Health and Human Services policies, and local and federal laws regarding animal welfare.

### Statistics

Significance was evaluated by the following analyses: student’s t-test for 2 groups, non-parametric Wilcoxon Rank Sum (Mann-Whitney) or Kruskal-Wallis for 3 or more groups, or Chi-square test for proportion comparison. Specific tests are denoted in the figure legends. Analyses were made in R using ggplot2 (3.5.1) and ggpubr (0.6.0) using stat_compare_means() and stat_summary() functions. *P* values were adjusted for multiple comparisons with the Bonferroni correction method when appropriate. Data are presented as means ± standard error (SEM) or means ± standard deviation (STDV). Individual data points are shown, or n is indicated. *P* < 0.05 was considered statistically significant.

### Data availability

Datasets supporting this study are available from corresponding author Dr. Christina V. Angeles upon request. Raw and processed DNA and RNA data will be made available through the National Institutes of Health Gene Expression Omnibus (GEO) database and dbGAP upon publication of the manuscript. Material or data that require a Material Transfer Agreement (MTA) can be provided by U-M pending scientific review and the execution of an MTA negotiated by the university’s Office of Technology Transfer. Requests for data that require an MTA should be submitted to the corresponding author.

## Author contributions

CVA conceptualized the study, designed experiments, interpreted data, wrote and finalized the manuscript, and supervised the project. AMS, EK, and LFG performed experiments, analyzed and interpreted data, and wrote and finalized the manuscript. AME and MKI performed bioinformatics analyses and finalized the manuscript. KDP and SCB performed pathological analysis, interpreted data, provided relevant advice, and finalized the manuscript. JW, WY, BK, JM, JMD, and CEE conducted experiments. CJG analyzed MRI images. WJ assisted with initial experimental design. AHC interpreted data and provided relevant advice. RC, TLF, MPM and AAD provided key materials, interpreted data and provided relevant advice. The co-first authors are listed according to the order they entered they study. All authors reviewed the manuscript.

## Acknowledgments

The authors thank our pathology technician, Deb Postiff, for her dedication and time procuring human tumors. We thank Drs. Ormond MacDougald and Kenneth Lewis for supporting and guiding adipocyte molecular biology techniques and confocal microscopy. We thank Lee Olsen for assistance with editing the manuscript. This work was supported by the National Cancer Institute (NCI) of NIH (R03CA280126 and NCI Cancer Center Support Grant P30CA046592 to CVA) and a Rogel Cancer Center Discovery Grant to CVA. CVA is also supported in part by the National Institute of Biomedical Imaging and Bioengineering of NIH Grant R01EB034399. LFG is supported by the National Institute of General Medical Sciences of NIH through the Training Program in Translational Research (T32GM141840). EK is supported by the University of Michigan (U-M) Doctoral Program in Cancer Biology. We thank the Rogel Cancer Center Tissue and Molecular Pathology, Flow Cytometry, Single Cell and Spatial Analysis, and Preclinical Molecular Imaging Shared Resources, which are supported by NCI P30CA046592; the Vector Core at the U-M Medical School; the U-M Advanced Genomics Core, supported by NCI P30CA046592, where library prep and next-generation sequencing was completed; and the Unit for Laboratory Animal Medicine of the U-M Medical School Office of Research. The authors thank the sarcoma patients whose tissue donations make this impactful research possible.

