## Supplemental Methods, Tables, Figures for "Immunocompetent murine models recapitulate the heterogeneous tumor immune microenvironment of human liposarcoma"

[1]Department of Surgery, [2]Program in Cancer Biology, [3]Department of Computational Medicine and Bioinformatics, [4]Department of Pathology, [5]Department of Dermatology, [6]Rogel Cancer Center, [7]Department of Pharmacology, [8]Department of Radiology, [9]Department of Internal Medicine, [10]Michigan Center for Translational Pathology, and [11]Department of Cell and Developmental Biology, University of Michigan, Ann Arbor, MI, 48109

***Authorship note:** AMS, EK, LFG contributed equally to this work and are co-first authors.

**Corresponding Author:**

Christina V. Angeles, MD, FACS, FSSO

University of Michigan, Rogel Cancer Center

1500 E. Medical Center Drive, 3304 Cancer Center

Ann Arbor, MI 48109

734-615-4823

Conflict of Interest: C.V.A. receives research funding (to the institution) from Skyline DX. K.D.P receives publishing royalties from Elsevier. The authors declare no relevant competing interests.

**Supplemental Methods**

*mT/mG indicator mice:* B6.129(Cg)-*Gt(ROSA)26Sor*$^{tm4(ACTB-tdTomato,-EGFP)Luo}$/J mice (Jackson Laboratory, stock #007676) were used to assess induction of the AAV8-Ap2.2-eGFP/Cre virus. Tissues were harvested 8 days post-injection for staining.

*Tumor monitoring and tissue harvesting.* Tumor growth was monitored 3 times per week using digital calipers. Mice were euthanized and all tissues were harvested when a flank tumor reached 20 mm in diameter, as per U-M Institutional Animal Care and Use Committee (IACUC) protocol. Tissues collected included tumors, spleens, lungs, liver and contralateral fat pad (when applicable). Fresh tissues were divided and stored in formalin fixed paraffin embedded (FFPE) blocks or in liquid nitrogen for future DNA and RNA extraction. Lungs, livers, and abdominal cavities were examined for macro-metastases at time of harvest. Lungs and livers were checked for microscopic disease via H&E. For virus-injected mice, if no tumors were observed 18 months following injection, mice were euthanized via $CO_2$ inhalation. For tail vein injections, mice were euthanized for tissue collection if a tumor started to develop in an undesired location or 6-7 months post injection.

*Magnetic Resonance Imaging (MRI) of ACPP mice.* Mice injected with AAV8-Ap2.2-eGFP/Cre were imaged via MRI every 2 - 4 weeks following injection. Following anesthetization with 2% isoflurane, mice were laid prone, headfirst, in a horizontal bore 7.0T Bruker MR scanner with body temperature maintained at 37°C using circulating heated water. A quadrature volume radiofrequency coil was used to scan the abdominal region. Gated coronal images were acquired using a T2-weighted rapid acquisition with relaxation enhancement (T2-TurboRARE) sequence with the following parameters: repetition time (TR)/effective echo time (TE), 3500/26 ms; echo spacing, 8.667 ms; number of echoes, 8; field of view (FOV), 40×50 mm; matrix, 256×128; slice thickness, 0.5 mm; number of slices, 40; and number of scans, 1.

*3T3 cell line differentiation and culturing.* CL-173 3T3-L1 (3T3) cells were purchased from ATCC (Lot #70043674). Cells were expanded using expansion cell media [DMEM (11965-084) with 10% CALF serum (ATCC 30-2030), 100 U/mL Penicillin/Streptomycin, 2 mM L-glutamine, 2.5 µg/mL plasmoscin]. Cells were cultured at 37°C in a humidified incubator with 10% $CO_2$. Cells were passaged and expanded when they approached confluency. Cells were differentiated using differentiation media [DMEM (11965-084) with 10% FBS, 100 U/mL Penicillin/Streptomycin, 250 mM IBMX (Sigma, I5879-250mg), 10 mg/mL insulin (Sigma, I0516-5mL), 10 mM Dexamethasone (Sigma, D4902-25mg), 25 mM Rosiglitazone (Sigma, R2408-10mg)] on days 1-3. Next, cells were cultured in high insulin media [DMEM (11965-084) with 10% FBS, 100 U/mL Penicillin/Streptomycin, 10 mg/mL insulin (Sigma, I0516-5mL)] on days 4-6. Media was replaced with expansion cell media on days 7-9 for final differentiation to lipid producing adipocytes.

*DNA and RNA Extraction***.** Tissue samples were stored in liquid nitrogen or RNAlater stabilization solution until extraction. 30 mg of frozen tissue was dissociated using the BeadBug Homogenizer (Southern Labware, D1030). DNA and RNA was extracted using the AllPrep DNA/RNA Mini Kit (Qiagen #80204). DNA and RNA concentrations were determined using the NanoDrop 2000 (Thermo Fisher Scientific).

*Quantitative reverse transcription PCR.* Samples were stored in liquid nitrogen or RNAlater and dissociated using the BeadBug Homogenizer. RNA was extracted using the RNeasy Mini Kit (Qiagen #74104). RNA was quantified using the NanoDrop 2000 and 200 ng was used for reverse transcription with the High-Capacity cDNA RT kit (Applied Biosystems #4368814). Cycle was 25°C 10 minutes, 37°C 2 hours, 85°C 5 minutes. qPCR was performed using Fast SYBR Green Master Mix (Thermo Fisher Scientific). Primers are listed in Supplemental Table 3. RNA levels were normalized to GAPDH RNA levels using the 2–ΔCt method.

*Whole Exome Sequencing (WES).* Whole exome sequencing was performed on genomic DNA from murine ACPP tumors (*n* = 14) on an Illumina NovaSeq 6000 by Novogene. Sample preparation, sample QC, library preparation, library QC, sequencing, and data QC were performed by Novogene. Exome enrichment was performed using Agilent SureSelectXT Mouse Exon Kit and 150 bp paired-end reads were generated with sequencing depth above 50X (6G).

*Bulk RNA sequencing.* Messenger RNA was purified from total RNA using poly-T oligo-attached magnetic beads. After fragmentation, first strand cDNA was synthesized using random hexamer primers, followed by second strand cDNA synthesis using dUTP for directional library or dTTP for non-directional library. The library was checked with Qubit and real-time PCR for quantification and bioanalyzer for size distribution detection. Quantified libraries were pooled and sequenced on Illumina platforms based on effective library concentration and data amount. Index-coded samples were clustered according to manufacturer instructions. After cluster generation, library preparations were sequenced on an Illumina platform and paired-end reads were generated. Raw reads in fastq format were processed through fastp software. Clean reads were obtained by removing low-quality reads and reads containing adapters and poly-N from raw data. At the same time, Q20, Q30 and GC content within clean data were calculated. All downstream analyses were based on clean data. Reference genome (GRCm39) and gene model annotation files were accessed from https://www.ncbi.nlm.nih.gov/grc/mouse/data. The index of the reference genome was built and paired-end clean reads were aligned to the reference genome using Hisat2 v2.0.5. featureCounts v1.5.0-p3 was used to count the read numbers mapped to each gene. Fragments per kilobase of transcript of each gene was calculated based on the length of the gene and reads counts mapped to this gene. Sequencing was performed by Novogene and the U-M Advanced Genomics Core.

*Spectral Flow Cytometry Sample Preparation.* Cell line-derived tumor samples were minced and digested in RPMI containing 2 mg/mL collagenase Type IV (Sigma, C5138-500mg) and 1 mg/mL DNase I (Sigma, 10104159001) at 37°C for 30 minutes. Dissociated cells were filtered through a 70 μM nylon mesh filter and washed with FACS buffer. Cells were then stained with the viability dye FVS780 for 30 minutes at 4°C. Upon removing the viability stain, Fc receptors were blocked with anti-mouse FC blocking Ab (Bio X Cell). Cells were then incubated for 45 minutes at 4°C for surface marker targets listed in Supplemental Table 2. Cells were washed with Flow buffer, fixed with Fixation Medium A for 15 minutes at room temperature and strained through a 35 μM filter cap tube and stored overnight at 4°C. Flow cytometry data were collected and compensated via unmixing on the Cytek Aurora at the U-M Flow Cytometry Core (Ann Arbor, Michigan). Data were analyzed using FlowJo™ v10 software (BD Life Sciences).

**Supplemental Table 1: Spontaneous Tumor Samples by Histology and Grade**

| Sample Number | Sex | Histology | Grade | TILs |
|---|---|---|---|---|
| N1006 | F | WDLPS | 1 | Low - Moderate |
| N1010 | F | WDLPS/DDLPS | 3 | Low-Moderate |
| N1019 | F | DDLPS | 3 | Low-Moderate |
| N1041 | F | WDLPS | 1 | Low |
| N1044 | F | WDLPS | 1 | Low |
| N1062 | F | WDLPS/DDLPS | 3 | Low-Moderate |
| N1104 | F | WDLPS | 1 | Low |
| N1107 | F | WDLPS | 1 | Low |
| N1108 | M | WDLPS/DDLPS | 3 | Low-Moderate |
| N1211 | M | WDLPS/DDLPS | 3 | Low-Moderate |
| N1216 | M | WDLPS/DDLPS | 3 | Low |
| N1255 | M | DDLPS | 2 | High |
| N1270 | M | WDLPS/DDLPS | 3 | High |
| N1285 | M | WDLPS/DDLPS | 3 | Low-Moderate |
| N1297 | M | DDLPS | 3 | Low-Moderate |
| N1300 | M | DDLPS | 3 | High |
| N1307 | F | WDLPS | 1 | Low |
| N1314 | F | WDLPS/DDLPS | 3 | High |
| N1319 | M | DDLPS | 3 | Low-Moderate |
| N1324 | M | WDLPS | 1 | Low |
| N1332 | M | DDLPS | 3 | Low-Moderate |
| N1339 | M | WDLPS/DDLPS | 2 | Low-Moderate |
| N1345 | F | WDLPS/DDLPS | 3 | High |
| N1350 | F | WDLPS | 1 | Low |
| N1359 | F | WDLPS/DDLPS | 3 | Low-Moderate |
| N1362 | F | WDLPS | 1 | Low |
| N1369 | F | DDLPS | 2 | High |

| | | | | |
|---|---|---|---|---|
| N1374 | F | WDLPS | 1 | Low |
| N1387 | F | WDLPS/DDLPS | 1 | Low |
| N1392 | F | WDLPS/DDLPS | 3 | Low-Moderate |
| N1396 | F | WDLPS | 1 | Low |
| N1429 | M | DDLPS | 3 | Low-Moderate |
| N1432 | M | WDLPS | 1 | Low |
| N1436 | F | WDLPS | 1 | Low |
| N1438 | F | WDLPS | 1 | Low |
| N1444 | M | WDLPS | 1 | Low |
| N1472 | M | WDLPS/DDLPS | 3 | Low-Moderate |
| N1474 | M | WDLPS | 1 | Low |
| N1508 | M | WDLPS | 1 | Low |
| N1514 | M | WDLPS | 1 | Low |
| N1518 | M | WDLPS/DDLPS | 2 | Low |
| N1528 | M | DDLPS | 3 | Low-Moderate |
| N1538 | F | WDLPS/DDLPS | 2 | Low-Moderate |
| N1541 | F | WDLPS | 1 | Low |
| N1549 | M | WDLPS | 1 | Low |
| N1551 | M | WDLPS/DDLPS | 2 | Low-Moderate |
| N1557 | M | WDLPS | 1 | Low |
| N1559 | M | WDLPS/DDLPS | 3 | Low-Moderate |
| N1564 | M | WDLPS | 1 | Low |
| N1567 | M | WDLPS/DDLPS | 3 | Low-Moderate |
| N1572 | M | DDLPS | 2 | Low |
| N1575 | M | WDLPS | 1 | Low |
| N1602 | F | WDLPS/DDLPS | 2 | Low-Moderate |
| N1607 | F | WDLPS | 1 | Low |
| N1621 | F | WDLPS/DDLPS | 3 | Low |
| N1624 | M | DDLPS | 3 | Low-Moderate |

| | | | | |
|---|---|---|---|---|
| N1629 | M | WDLPS | 1 | Low-Moderate |
| N1637 | F | WDLPS | 1 | Low-Moderate |
| N1638 | F | WDLPS | 1 | Low |
| N1661 | F | WDLPS | 1 | Low |
| N1667 | F | WDLPS/DDLPS | 2 | Low-Moderate |
| N1690* | F | WDLPS/DDLPS (WD part) | 3 | Low-Moderate |
| N1692* | F | WDLPS/DDLPS (DD part) | 3 | Low-Moderate |
| N1706 | M | WDLPS | 1 | Low |
| N1711 | M | DDLPS | 3 | Low |
| N1715 | F | WDLPS | 1 | Low |
| N1723 | M | Cellular WDLPS | 1 | Low-Moderate |
| N1733 | M | WDLPS | 1 | Low |
| N1770 | F | WDLPS/DDLPS | 3 | Low-Moderate |
| N1783 | F | WDLPS | 1 | Low |
| N1785 | F | DDLPS | 3 | Low |
| N1791 | F | DDLPS | 2 | High |
| N1837 | M | WDLPS | 1 | Low |
| N1840 | M | DDLPS | 3 | Low-Moderate |
| N1852 | F | WDLPS | 1 | Low |
| N1855 | F | DDLPS | 3 | Low |
| N1867 | F | WDLPS | 1 | Low-Moderate |
| N1870 | F | WDLPS | 1 | Low |
| N1879 | F | WDLPS | 1 | Low |
| N1882 | F | WDLPS | 1 | Low |
| N1940 | F | WDLPS | 1 | Low-Moderate |

*Samples N1690 and N1692 came from the same spontaneous ACPP tumor, N1689.

**Supplemental Table 2: Antibodies**

| Antibodies | Supplier | Catalog # | IHC Dilution | IF Dilution |
|---|---|---|---|---|
| F480 | Cell Signaling Technology | 70076S | 1:500 | - |
| CD3 (Standard)** | Abcam | AB11089 | 1:100 | - |
| CD8α (Standard)** | Cell Signaling Technology | 98941S | 1:200 | - |
| CD19 | Invitrogen | PIPA527442 | 1:500 | - |
| PD1 | Abcam | AB214421 | 1:1000 | - |
| FOXP3 | Cell Signaling Technology | 12653S | 1:800 | - |
| CD3 ($T_{RM}$)** | Dako | A0452 | 1:100 | - |
| CD8α ($T_{RM}$)** | Abcam | AB178081 | 1:100 | - |
| CD103 | Abcam | AB129202 | 1:500 | - |
| CD69 | Abcam | AB233396 | 1:500 | - |
| Desmin | Abnova | MAB19978 | 1:3000 | - |
| GFP | Abcam | AB6673 | - | 1:200 |
| **Flow Cytometry Antibodies** | **Supplier** | **Fluorophore** | **Clone/Lot** | **Dilution** |
| CD45 | BD | BV510 | 30-F11 | 1:100 |
| CD3 | BioLegend | FITC | 17A2 | 1:100 |
| CD4 | BioLegend | BV570 | RM4-5 | 1:100 |
| CD8α | BioLegend | BV650 | 53-6.7 | 1:100 |
| CD69 | BioLegend | APC | H1.2F3 | 1:100 |
| CD103 | BD | PE-CF594 | M290 | 1:100 |
| CD19 | BioLegend | Pacific Blue | ID3/CD19 | 1:100 |
| PD1 | BD | APC-R700 | J43 | 1:100 |
| LAG3 | BioLegend | PE-Cy7 | C9B7W | 1:100 |
| CD44 | BD | BB700 | IM7 | 1:100 |
| CXCR6 (CD186) | BioLegend | BV711 | SA051D1 | 1:100 |
| CD11b | BioLegend | BV785 | M1/70 | 1:100 |
| F480 | BD | BV480 | T45-2342 | 1:100 |
| CD62L | BioLegend | BV605 | MEL-14 | 1:100 |
| Viability | Invitrogen | Fixable Viability Dye eFluor$^{TM}$ 780 | 2851407 | 1:1000 |

**Different CD3 and CD8α primary antibodies were used in the two mIHC panels (Standard and $T_{RM}$) due to fluorescent dye compatibility considerations during panel design.

### Supplemental Table 3: Primers

| Target Gene (murine) | Forward | Reverse |
|---|---|---|
| *Trp53* | TTT GAA GGC CCA AGT GAA GC | CTG ACC CAC AAC TGC ACA G |
| *Pten* | GAA AGG GAC GGA CTG GTG TA | AGT GCC ACG GGT CTG TAA TC |
| *Gapdh* | CAT CAC TGC CAC CCA GAA GAC TG | ATG CCA GTG AGC TTC CCG TTC AG |
| *Pparg* | GTA CTG TCG GTT TCA GAA GTG CC | ATC TCC GCC AAC AGC TTC TCC T |
| *Cebpα* | GCA AAG CCA AGA AGT CGG TGG A | CCT TCT GTT GCG TCT CCA CGT T |
| *AdipoQ* | AGA TGG CAC TCC TGG AGA GAA G | ACA TAA GCG GCT TCT CCA GGC T |
| *Fabp4* | TGA AAT CAC CGC AGA CGA CAG G | GCT TGT CAC CAT CTC GTT TTC TC |

### Supplemental Figures

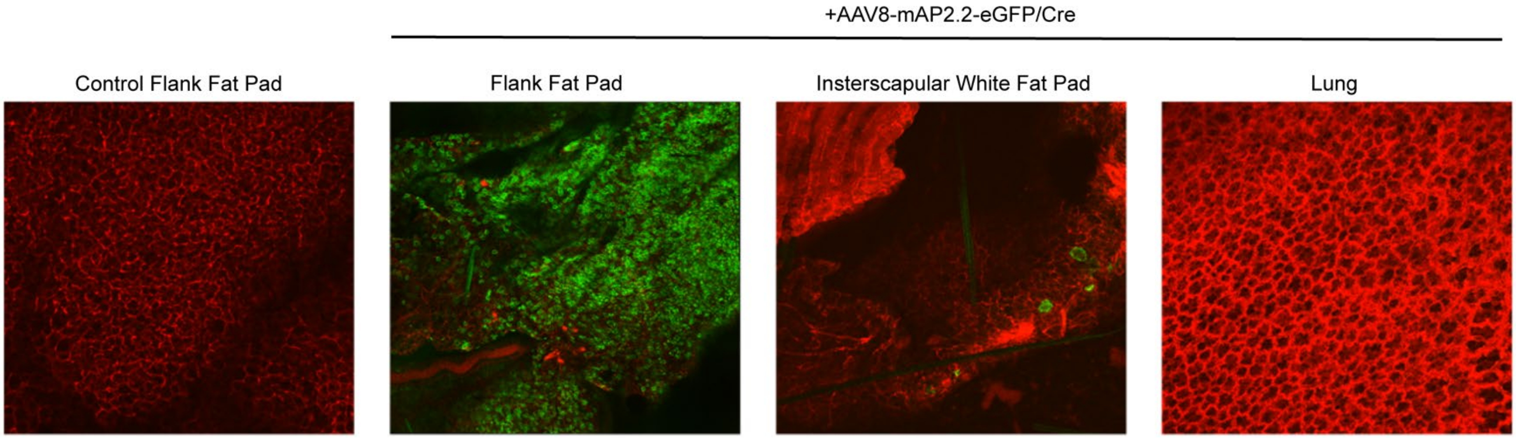

**Supplemental Figure 1. Validation of adipocyte-specific transduction and recombination using AAV8-mAP2.2-eGFP/Cre in mT/mG mice.** mT/mG mice show ubiquitous tdTomato (red) expression in adipose tissue. Injection of $1 \times 10^{11}$ vg AAV8-mAP2.2-eGFP/Cre into flank fat pad shows successful Cre-mediated recombination, loss of tdTomato, and activation of GFP expression (green). Distant interscapular fat has minimal recombination (GFP expression), and lung shows no detectable recombination at this distant site.

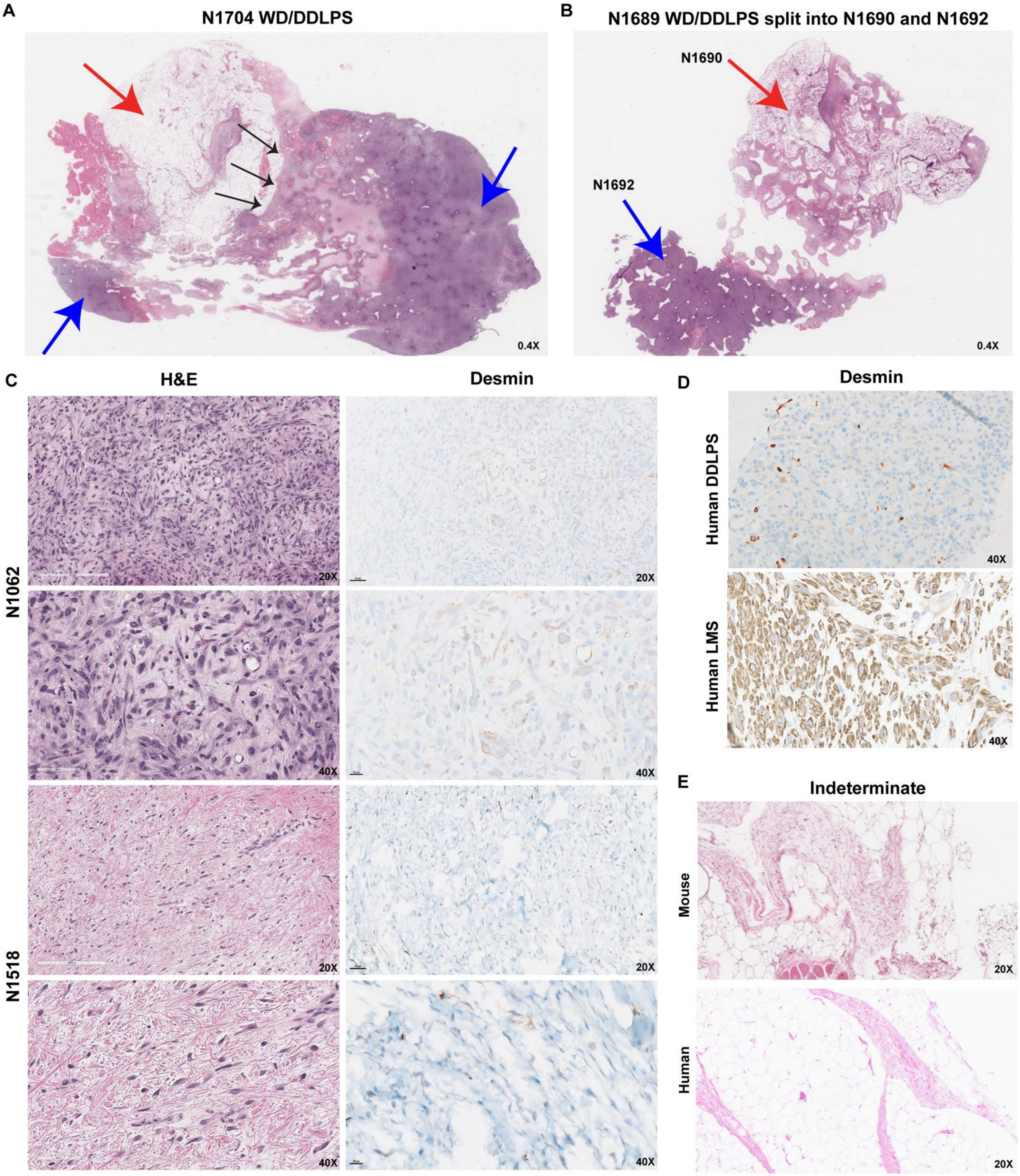

**Supplemental Figure 2. Autochthonous ACPP tumors exhibit human LPS histological subtypes.** (**A-B**) Representative H&Es of ACPP tumors with both WDLPS and DDLPS regions represented by red and blue arrows, respectively (0.4X). Black arrows indicate region of WD-DD transition. (**C**) Representative H&E and desmin chromogenic IHC staining for ACPP tumors (*n* = 2, 20X and 40X). Scale bar shown. (**D**) Representative human DDLPS (negative) and human LMS (positive) desmin chromogenic IHC (40X). (**E**) Example H&Es of murine ACPP tumor and human lipomatous tumor with histological pathological classification of indeterminate (20X).

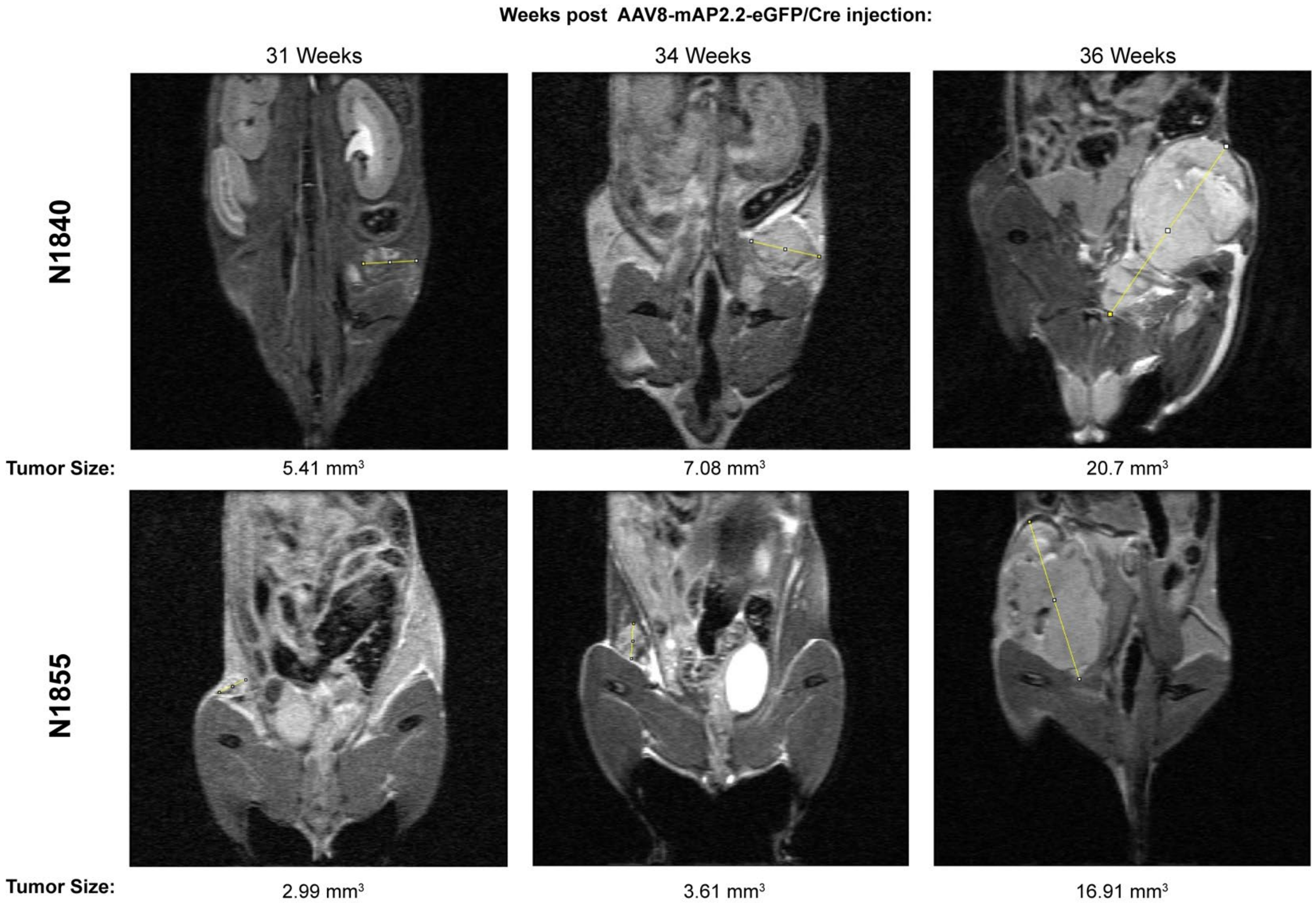

**Supplemental Figure 3. MRI enables early detection and progressive size tracking of ACPP tumors.** Sequential MRI images of two representative ACPP mice with autochthonous DDLPS. MRI enabled detection of ACPP tumors prior to detected by manual palpation. Tumor volume ($mm^3$) was calculated from MRI images using Voxel size equation.

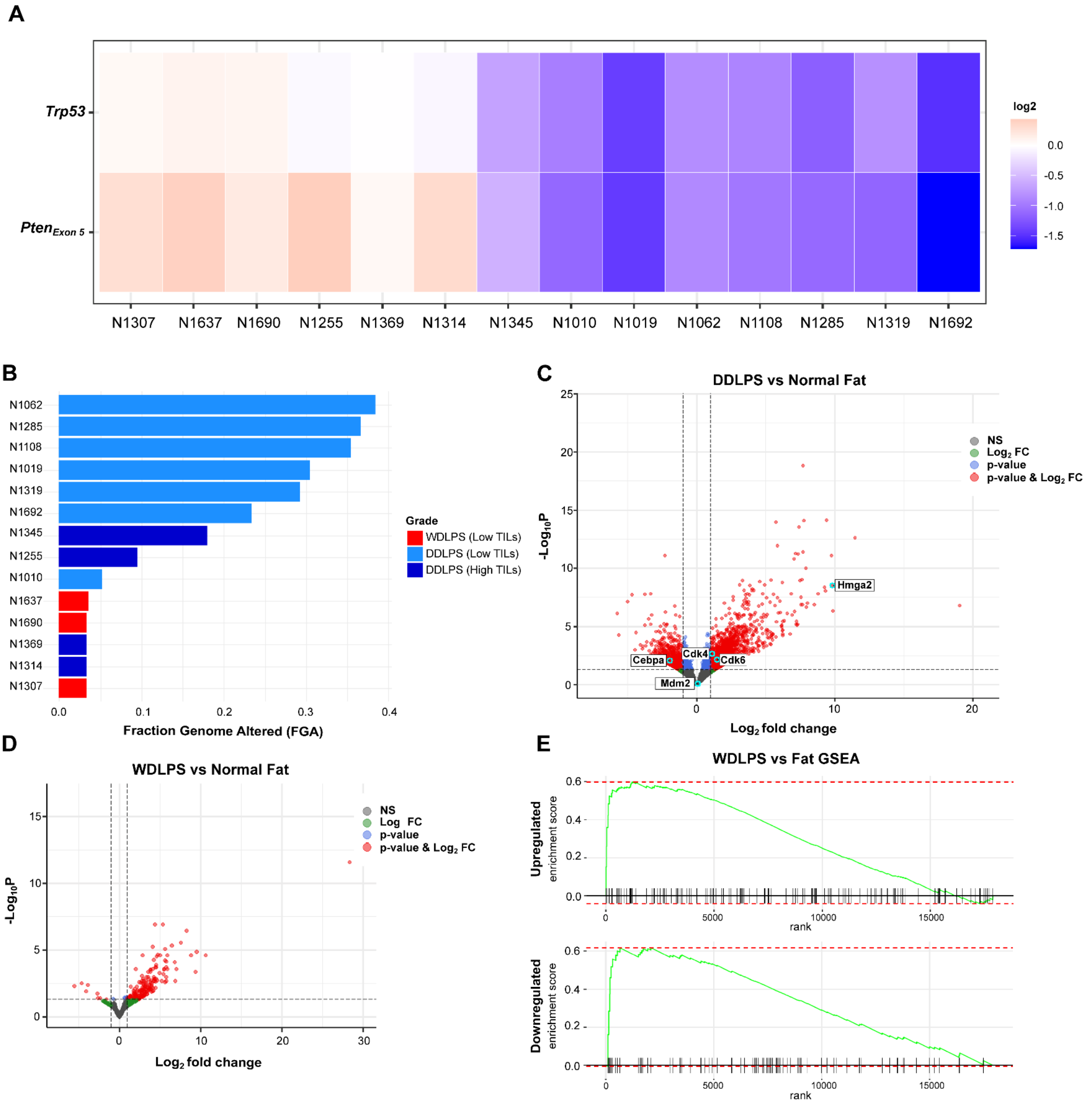

**Supplemental Figure 4. Multi-omic analysis of ACPP tumors.** (**A**) Heatmap of WES illustrating the loss of *Trp53* and *Pten* across ACPP tumor samples (*n* = 14) from AAV8-Ap2.2/Cre Trp53$^{f/f}$Pten$^{f/f}$ C57BL/6. (**B**) Bar graph of Fraction of Genome Altered (FGA) values for each ACPP tumor (*n* = 14); color corresponds to specified histological classification. (**C**) Volcano plot of differential gene expression (DGE) in murine DDLPS (*n* = 8) and (**D**) in murine WDLPS (*n* = 3) in comparison to normal fat. Red dots represent statistically significant downregulated and upregulated genes on the left and right side of zero, respectively. Threshold for significance was set at $P < |0.05|$ and $Log_2$ fold change > |0.5|. Genes of note are labeled. (**E**) GSEA enrichment plots between murine and human WDLPS for up- and down-regulated genes compared to species-specific normal fat.

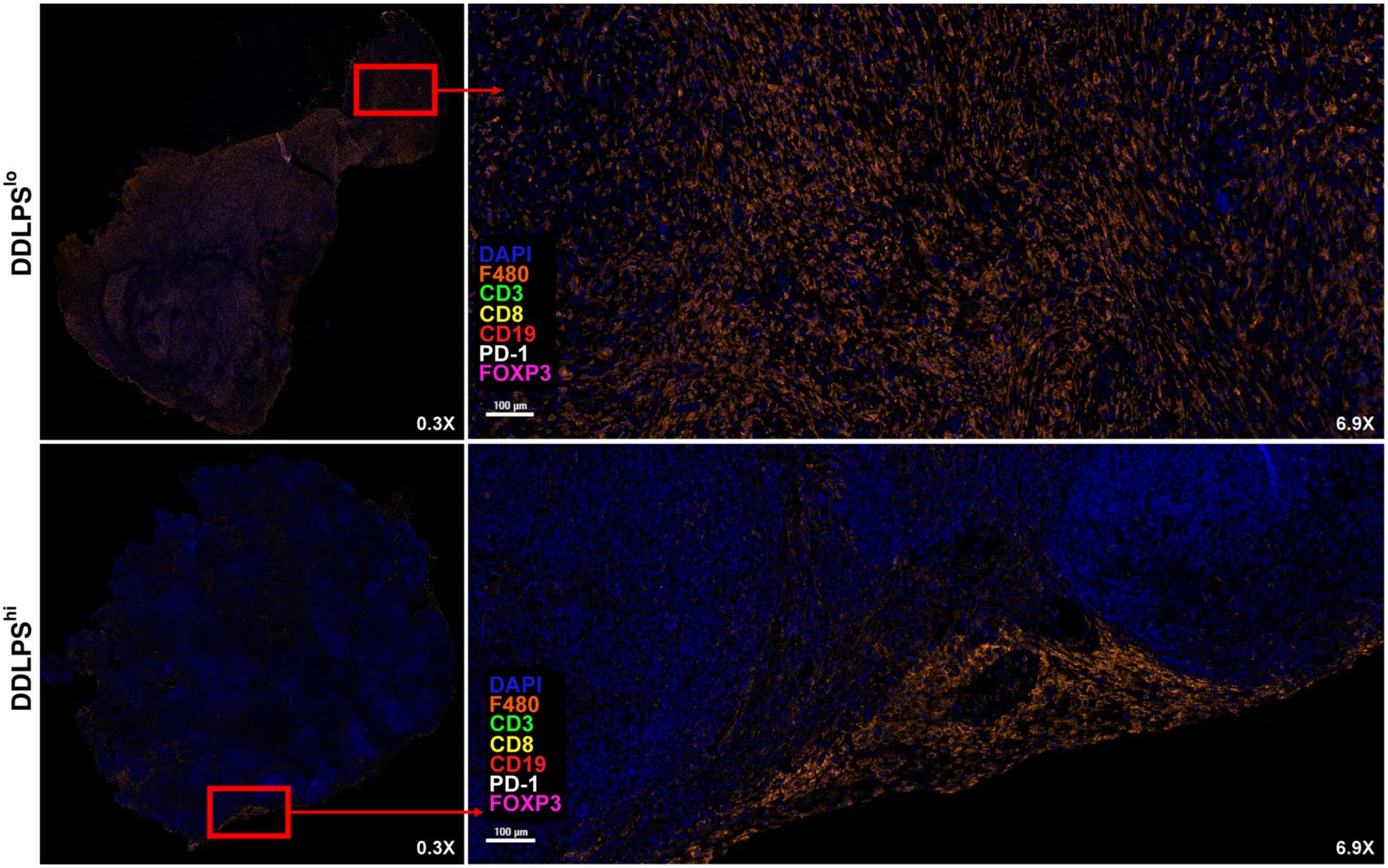

**Supplemental Figure 5. Macrophages have distinct spatial patterns between DDLPS TIME subtypes.** Multiplex IHC (mIHC) of representative ACPP DDLPS$^{lo}$ and DDLPS$^{hi}$ tumors with standard immune panel. DDLPS$^{hi}$ were defined as those with $CD3^{+}$ T cells ≥ 2x the median across all samples; DDLPS$^{lo}$ tumors had $CD3^{+}$ levels below this threshold. mIHC visualizes discrete spatial patterns of F480+ macrophages (orange) in DDLPS$^{lo}$ versus DDLPS$^{hi}$. DAPI (Hoechst/blue) used as a counterstain. Red boxes and arrows indicate area shown at higher magnifications (0.3X, 6.9X).

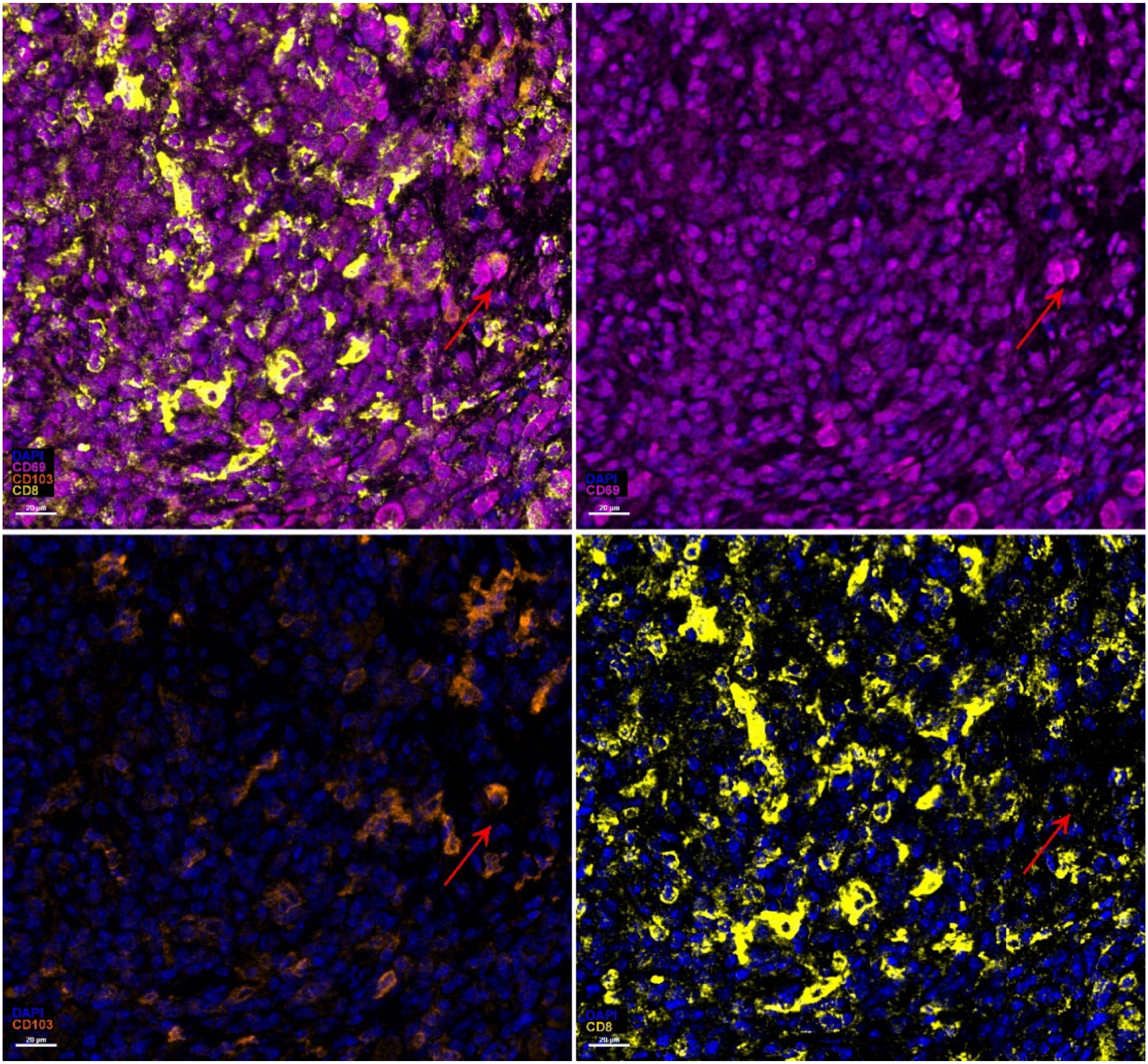

**Supplemental Figure 6. CD69$^+$CD103$^+$ CD8$^+$ resident memory ($T_{RM}$) cells are identified by mIHC.** Multiplex IHC (mIHC) of representative ACPP DDLPS$^{hi}$ tumor. mIHC $T_{RM}$ panel reveals T cells with colocalization of surface markers CD103 (orange) and CD69 (magenta) with DAPI (Hoechst) as a counterstain. Red arrows indicate identified cell with overlapping fluorescence. Shown at 40X magnification. Scale bar shown.

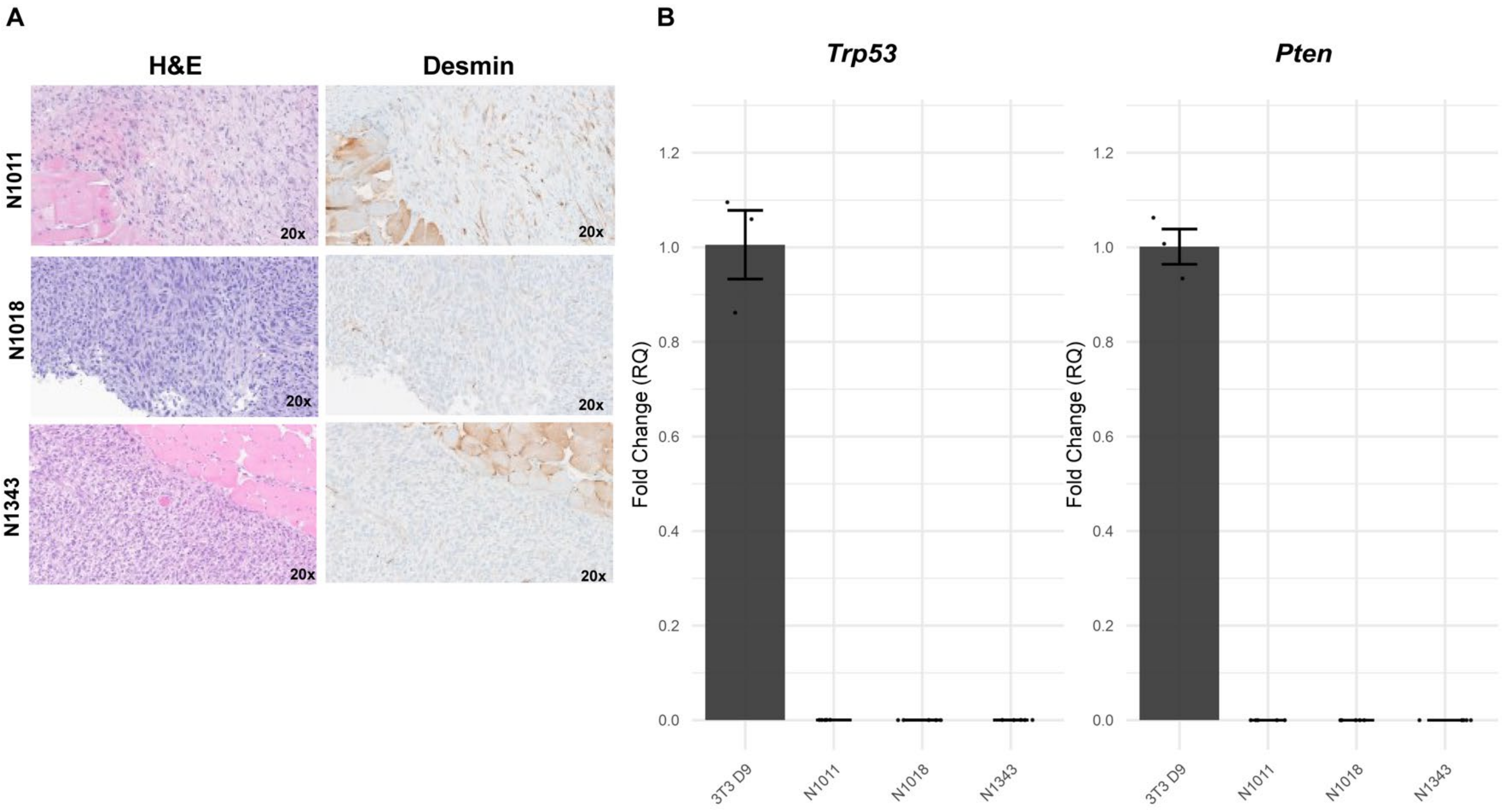

C

| ACPP Tumor | Sex | Histology | Grade | TILs (A) | Syngeneic Cell Line | Histology | Grade | TILs (B) | Mets |
|---|---|---|---|---|---|---|---|---|---|
| N1010 | F | DDLPS | 3 | Moderate | N1011 | DDLPS | 3 | + | No |
| N1019 | F | DDLPS | 3 | Moderate | N1018 | DDLPS | 3 | ++ | Yes |
| N1345 | F | DDLPS | 3 | High | N1343 | DDLPS | 3 | +++ | No |

**Supplemental Figure 7. Syngeneic cell line allografts retain characteristics and gene expression patterns observed in autochthonous ACPP tumors.** (**A**) Representative H&E and desmin chromogenic IHC staining for syngeneic allografts from each cell line (N1011, N1018, and N1343). (**B**) Analysis of *Trp53* and *Pten* gene expression levels via quantitative PCR (qPCR) in mature adipocytes (pre-adipocytes treated with differentiation factors to reach mature differentiation at day 9 = 3T3 D9) and our syngeneic cell lines (N1011, N1018, N1343). (**C**) Table of characteristics of each syngeneic cell line allograft and their corresponding originating ACPP tumor.

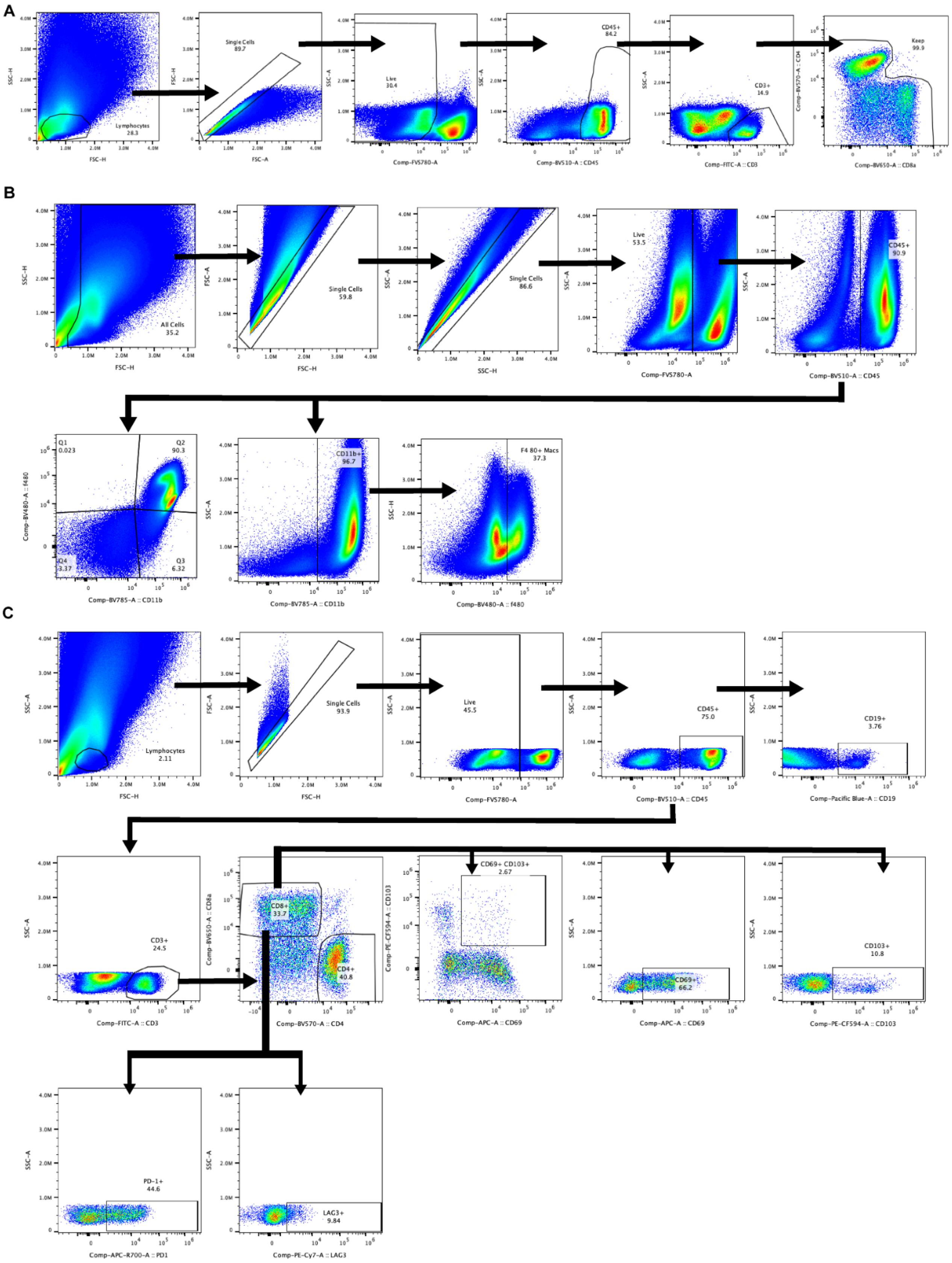

**Supplemental Figure 8. Gating strategies for spectral flow cytometry analysis.** (**A**) Pre-gating strategy for $CD3^{+}$ t-SNE clustering: lymphocytes/single cells/live/$CD45^{+}$/$CD3^{+}$. Gating was stringent for live, $CD45^{+}$ and $CD3^{+}$ to ensure clean data for clustering. Additional "keep" gate was used to remove $CD4^{+}$/$CD8^{+}$ double positive cells. (**B**) Myeloid populations gating strategy: all cells/single cells/live/$CD45^{+}$/$CD11b^{+}$ for myeloid cells; all cells/single cells/single cells/live/$CD45^{+}$/$CD11b^{+}F480^{+}$ for $CD11b^{+}$/$F480^{+}$ macrophages; all cells/single cells/live/$CD45^{+}$/$CD11b^{+}$/$F480^{hi}$ for $F480^{hi}$ macrophages. (**C**) Lymphocyte populations gating strategy: lymphocytes/single cells/live/$CD45^{+}$/$CD19^{+}$ for B cells; lymphocytes/single cells/live/$CD45^{+}$/$CD3^{+}$ for T cells; lymphocytes/single cells/live/$CD45^{+}$/$CD3^{+}$/$CD8^{+}$ or $CD4^{+}$ for $CD8^{+}$ T cells or $CD4^{+}$ T cells. All other markers were pre-gated on lymphocytes/single cells/live/$CD45^{+}$/$CD3^{+}$/$CD8^{+}$ for further characterization.

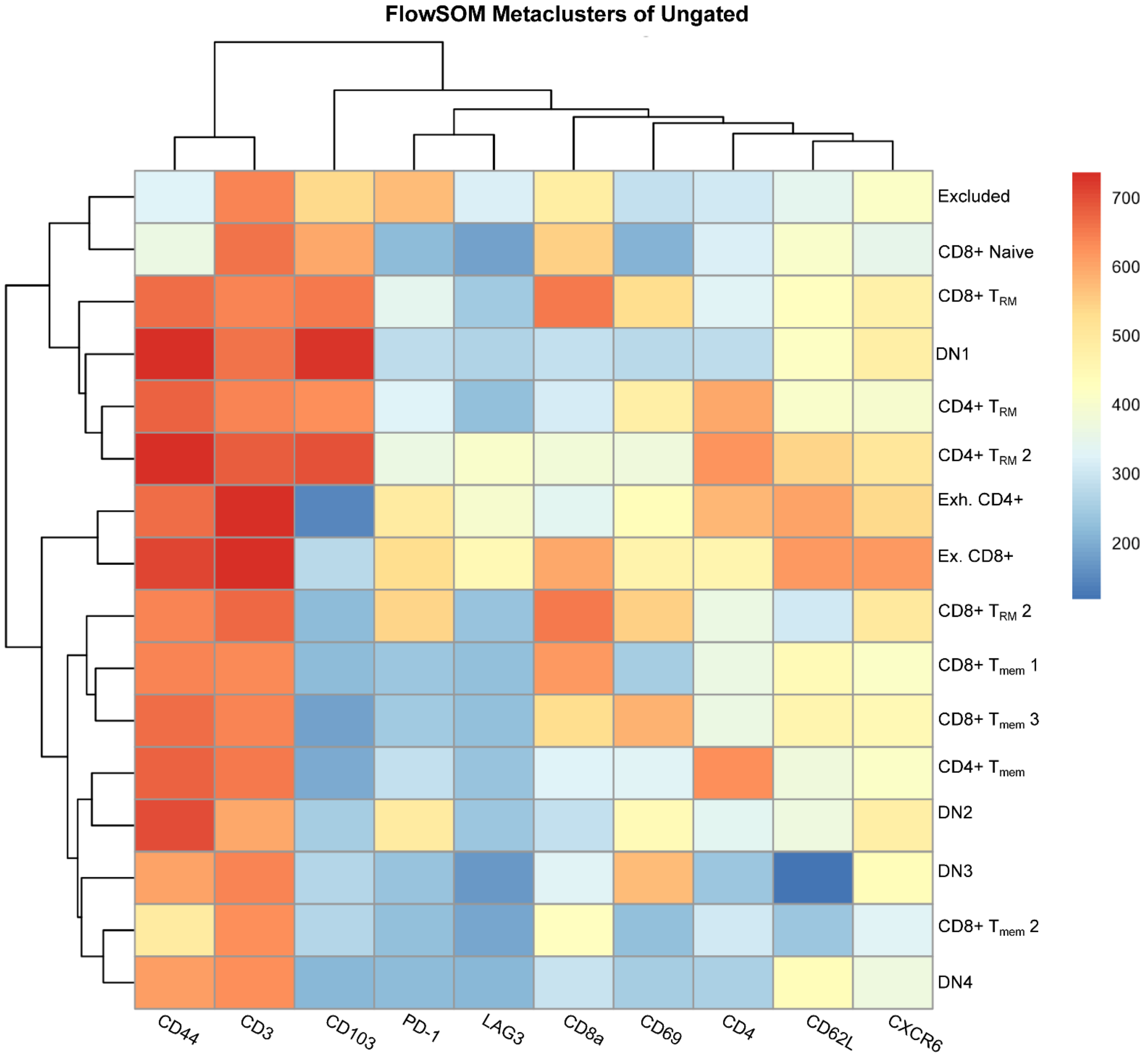

**Supplemental Figure 9. Heat map of FlowSOM CD3+ T cell meta-clusters.** Hierarchical heat map of marker expression via mean fluorescent intensities (MFI) among FlowSOM identified meta-clusters, which were used to name our CD3$^+$ T cell populations. Scale bar of MFI is shown.

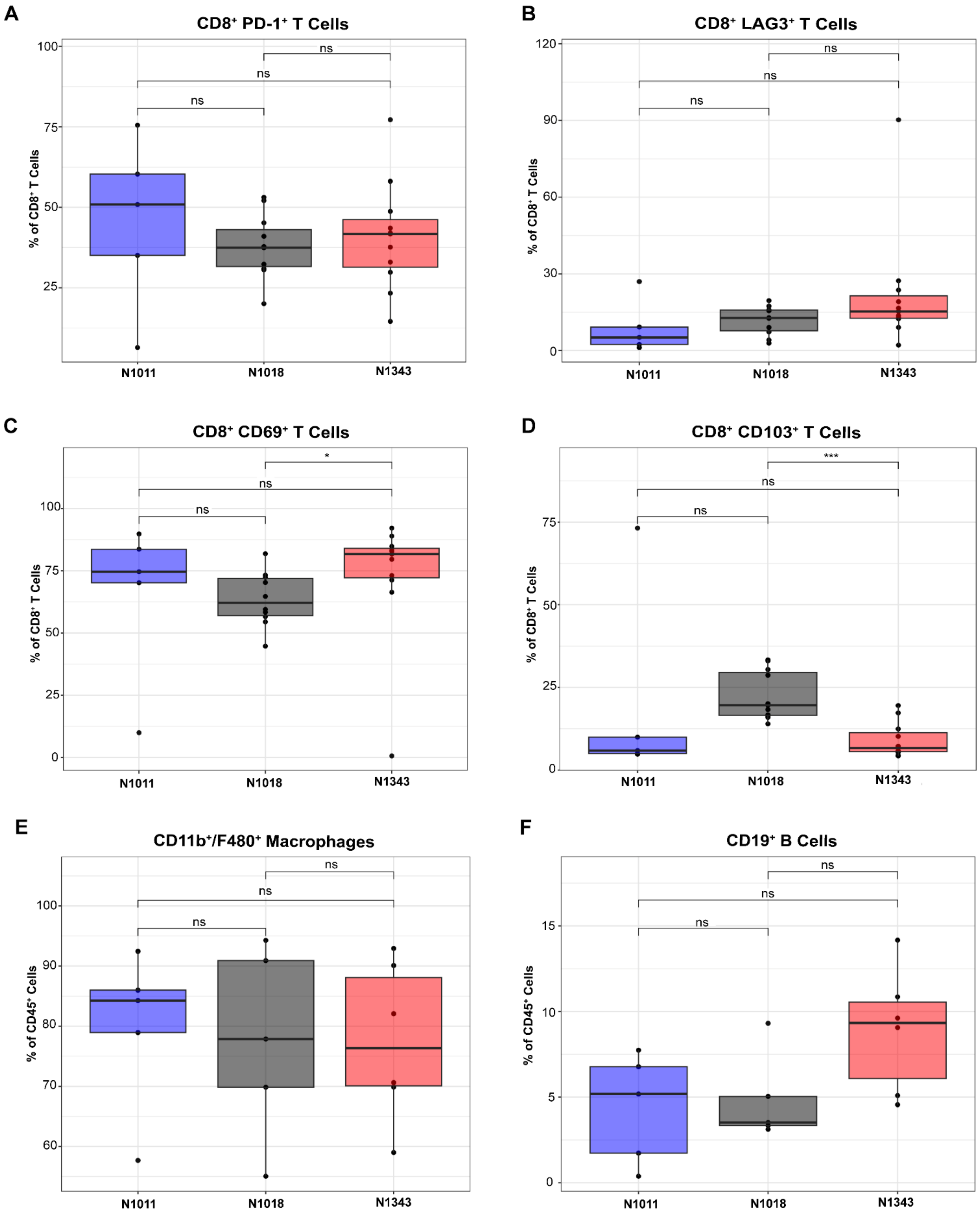

**Supplemental Figure 10. Spectral flow cytometry analysis of immune populations in syngeneic orthotopic allografts.** Comparisons of the percentages of CD8+ T cells with markers for (**A**) PD-1$^+$, (**B**) LAG3$^+$, (**C**) CD69$^+$ and (**D**) CD103$^+$, among syngeneic orthotopic allografts (N1011, N1018, or N1343 implanted in WT C57BL/6 mice). Pre-gated on lymphocytes/singlets/live/CD45$^+$ ($n$ = 5 – 10 per cell line; see gating strategy in Supplemental Figure 7B). (**E**) Percentage of CD45$^+$ cells that are CD11b$^+$/F480$^+$ macrophages among syngeneic allografts. Pre-gated on all cells/singlets/live/CD45$^+$ ($n$ = 5 - 10 per cell line; see gating strategy in Supplemental Figure 7C). (**F**) Percentage of CD45$^+$ cells that are CD19$^+$ B cells among syngeneic allografts. Pre-gated on lymphocytes/singlets/live/CD45$^+$. The box plots represent mean (center line), the length of the box represents the 25th to 75th percentiles and the whiskers represent the 5th to 95th percentiles of the data. Outliers are shown as dots outside of whiskers. P values were calculated by the Kruskal-Wallis test. Significance is indicated by * = $P < 0.05$ ; *** = $P < 0.001$; **** = $P < 0.0001$.
